# Efficient generation of transgene-free canker-resistant *Citrus sinensis* using Cas12a/crRNA ribonucleoprotein

**DOI:** 10.1101/2023.02.12.528187

**Authors:** Hang Su, Yuanchun Wang, Jin Xu, Ahmad A. Omar, Jude W. Grosser, Milica Calovic, Liyang Zhang, Christopher A. Vakulskas, Nian Wang

## Abstract

Citrus is one of the top three tree crops in the world and its production faces many devastating disease challenges such as citrus canker caused by *Xanthomonas citri* subsp. citri (Xcc). Genetic improvement of citrus via traditional approaches is a lengthy (approximately 20 years) and laborious process. Biotechnological approaches including CRISPR genome editing technologies have shown promise. However, none of the citrus plants generated by biotechnological approaches have been commercialized, which primarily resulted from the transgenic nature of the genetically modified plants. Here, we successfully developed transgene-free canker-resistant *Citrus sinensis* lines in the T0 generation within 10 months through transformation of embryogenic citrus protoplasts with Cas12a/crRNA ribonucleoprotein (RNP) to edit the canker susceptibility gene *CsLOB1*. Among the 39 regenerated lines, 38 are biallelic/homozygous mutants based on Sanger sequencing analysis of targeting sites and whole genome sequencing, demonstrating a 97.4% biallelic/homozygous mutation rate. The edited lines do not contain off-target mutations. The *CsLOB1* edited *C. sinensis* lines demonstrate no differences from wild type plants except canker resistance. Importantly, the transgene-free canker-resistant *C. sinensis* lines have received regulatory approval by USDA APHIS. This study presents an efficient genome editing approach for citrus using Cas12a/crRNA RNP, which has a broad impact on genetic improvement of elite citrus varieties and potentially other tree crops and their genetic study. This study represents a breakthrough by generating the first transgene-free canker-resistant *C. sinensis* lines that provide a sustainable and efficient solution to control citrus canker.

---

Citrus canker caused by *Xanthomonas citri* subsp. citri (Xcc) causes severe yield, quality and economic loss to citrus production worldwide and is endemic in most citrus-producing countries such as U.S., Brazil, and China ^1^. Xcc encodes a dominant pathogenicity factor PthA4^2,3^, a transcription activator-like effector (TALE) secreted by the type III secretion system. PthA4 enters the nucleus of plant cells to activate the canker susceptibility gene *LOB1* by binding to the effector binding elements in its promoter region, which subsequently induces expression of downstream genes and causes typical canker symptoms including hypertrophy and hyperplasia ^3^. All commercial citrus cultivars are susceptible to citrus canker ^4,5^. Citrus canker control relies primarily on treatment with copper-based antimicrobials ^6^, which cause environmental pollution ^7–9^. Furthermore, copper-resistant Xcc strains have been reported in citrus producing regions where copper was frequently used to control citrus canker ^10,11^. Numerous citrus breeding programs worldwide have been aiming to improve canker resistance for elite citrus cultivars. Traditional breeding to generate canker-resistant citrus varieties has been hindered by heterozygosity, long juvenile period, self- and cross-incompatibility. The average breeding duration from the cross to the release of a cultivar for traditional citrus breeding requires approximately 20 years ^12^. Transgenic expression of antimicrobial peptides ^13–15^, toxin ^16^, resistance genes ^17,18^, and immune-related genes ^19–25^ enabled increased canker resistance. Recently, CRISPR/Cas-mediated genome editing of the promoter or coding region of *LOB1* has conferred citrus resistance to Xcc ^26–32^. However, the citrus plants generated by transgenic overexpression and CRISPR genome editing approaches were all transgenic. Transgenic crops face many challenges to be used in production owing to regulations and public perception concerns ^33,34^. Consequently, none of the citrus plants generated by biotechnological approaches have been registered and commercialized despite the tremendous effort and superior disease resistance.

This study aimed to generate transgene-free canker-resistant *C. sinensis* varieties. Cas9 and Cas12a DNA, RNA or ribonucleoprotein complex (RNP) were successfully used to generate transgene-free crops in the T0 generation ^35,36^, which significantly shortens the time for plant genetic improvement by avoiding the lengthy process needed to remove transgenes.

Specifically, the Cas/gRNA RNP method does not involve DNA fragments and has been used to generate transgene-free tobacco, Arabidopsis, lettuce, rice ^36^, Petunia ^37^, grapevine, apple ^38^, maize ^39^, wheat ^40^, and potato ^41^. The RNP method is also known to reduce off-target mutations. For instance, off-target mutations were not detected for genome-edited maize ^39^ and wheat ^40^ that were generated by the RNP method. However, RNP-mediated genome editing efficacy is low ^42^. Here, we have generated transgene-free canker-resistant *C. sinensis* cv. Hamlin (a prevalent citrus cultivar) lines using LbCas12a/crRNA RNP, which do not contain off-target mutations, within 10 months (Fig. 1). Importantly, among the 39 regenerated lines, 38 lines were biallelic/homozygous mutants. The high efficacy and short time needed for Cas12a/crRNA RNP-mediated citrus genome editing will significantly impact how citrus and other tree crops are genetically improved as well as their genetic studies in the future.

**Figure 1.**
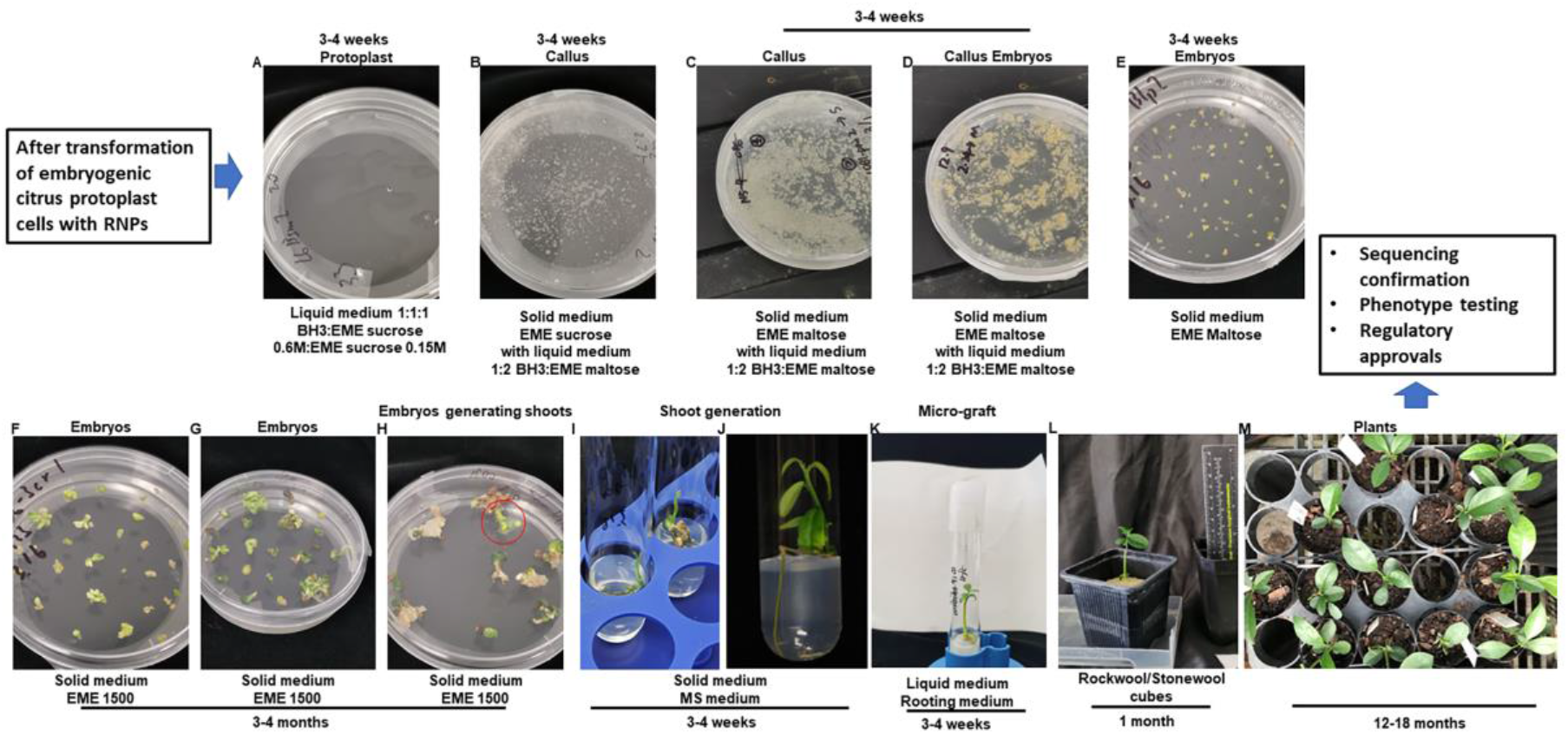
Regeneration of genome-edited citrus protoplast cells. A. Edited citrus protoplast cells were kept in liquid medium (1:1:1 (v:v:v) mixture of BH3 and EME sucrose 0.6 M and EME sucrose 0.15 M) for 3-4 weeks at 28°C in dark without shaking. B. Citrus cells were transferred to EME sucrose medium added with 1:2 mixture of BH3 and EME maltose 0.15 M and kept at 28°C for 3-4 weeks in dark. C and D. Calli were transferred to EME maltose solid medium added with 1:2 mixture of BH3 and EME maltose 0.15 M and kept at 28°C in dark for 3-4 weeks to generate embryos. E. Embryos were transferred to EME maltose solid medium and kept at room temperature under light for 3-4 weeks. F-H, Embryos were transferred to solid EME1500 medium and kept at room temperature under light for 3-4 months to generate shoots. I&J. Small plantlets were transferred to MS medium and kept at room temperature for 3-4 weeks. K-M. The regenerated shoots were micro-grafted onto Carrizo citrange rootstock in liquid rooting media and kept in tissue culture room at 25°C under light for 3-4 weeks (K), grown in stonewool cubes in growth chamber at 25°C under light for 1 month (L), then planted in soil (M).

## Evaluating the genome editing efficacy of LbCas12a/crRNA RNP and ErCas12a/crRNA RNP transformation of embryogenic citrus protoplasts by targeting the *CsPDS* gene

To evaluate the transgene-free citrus genome editing efficacy of the RNP method, we first used the *CsPDS* gene (orange1.1t02361) as the target owing to its obvious albino phenotype which expedited the identification of biallelic/homozygous mutations ^43^. Both Cas9 and Cas12a were successfully used in genome editing with the RNP method ^36,40,44^. Here, we selected Cas12a because it generates longer deletion than Cas9 ^26,45^. We assessed both ErCas12a and LbCas12a-Ultra (hereafter LbCas12aU) in the RNP-mediated genome editing of embryogenic citrus protoplasts. Both ErCas12a and LbCas12a were reported to have high genome editing efficacy ^46,47^. LbCas12aU is a variant of LbCas12a and has increased genome editing efficacy than LbCas12a. We first evaluated their efficiency via *in vitro* digestion of a 555 bp DNA fragment from the first exon of the *CsPDS* gene (Extended Data Fig. S1A). Both ErCas12a and LbCas12aU were able to digest the DNA fragments efficiently and generated two DNA fragments with the expected sizes of 320 bp and 240 bp (Extended Data Fig. S1B&C). LbCas12aU showed a slightly higher efficiency than ErCas12a in *in vitro* digestion (Extended Data Fig. S1B&C).

Next, LbCas12aU/crRNA and ErCas12a/crRNA RNPs were used to transform embryogenic *C. sinensis* cv. Hamlin protoplasts using the PEG method ^48^, which were used for plant regeneration without herbicide or antibiotics selection (Fig. 1). Six months after transformation with ErCas12a, 58 embryos showing an albino phenotype were selected for further analysis (Fig. 2a). Sanger sequencing analysis of the *CsPDS* gene indicated that the 58 albino mutants consisted of 56 homozygous mutants, 1 biallelic mutant, and 1 chimeric mutant. A randomly selected green embryo contained no mutations at the target site (Fig. 2). Among the homozygous mutants, 30 contained 7 bp deletion, whereas 26 contained 13 bp deletion. The biallelic mutant T0_Er_-22 contained both 7 bp and 13 bp deletions, and the chimeric mutant T0_Er_-20 contained 7 bp, 8 bp and 13 bp deletions (Fig. 2b). For LbCas12aU, 15 embryos showing an albino phenotype were selected for Sanger sequencing analyses (Fig. 2a). The sequencing result demonstrated that all the embryos generated from LbCas12aU/crRNA RNP transformation contained the same mutation type (9 bp deletion) at the target site (Fig 2b). Thus, we concluded that both LbCas12aU/crRNA and ErCas12a/crRNA RNP transformation of embryogenic citrus protoplasts were able to efficiently generate biallelic/homozygous *CsPDS* mutations for *C. sinensis*.

**Figure 2.**
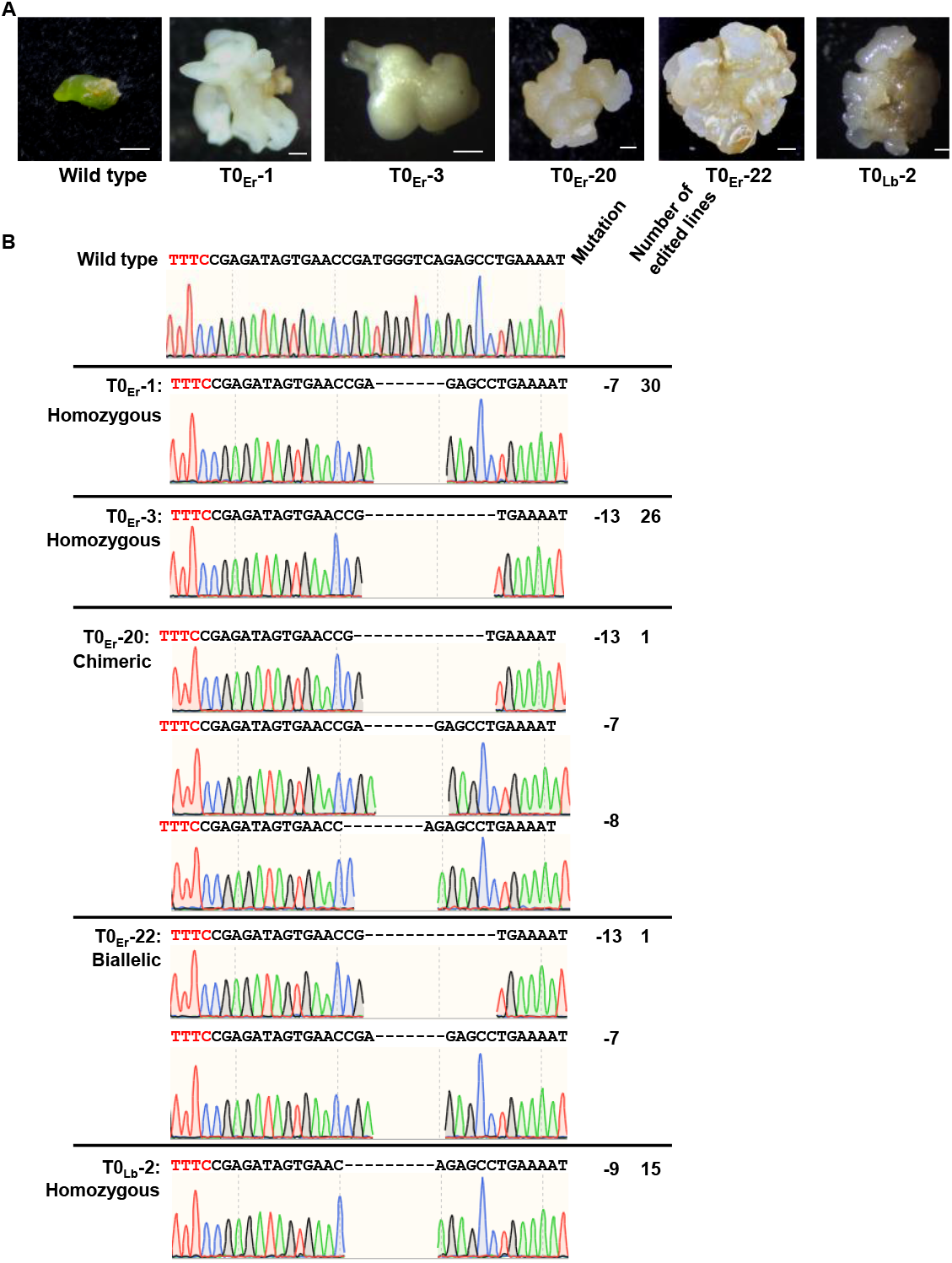
Genome editing of the *CsPDS* gene of embryogenic protoplast cells of *C. sinensis* cv. Hamlin. A. Genome-edited embryos under regeneration. Er indicates ErCas12a. Lb indicates LbCas12aU. B. Sanger sequencing confirmation of genome-edited lines in the *CsPDS* gene. TTTC in red: PAM (protospacer adjacent motif). Mutation indicates mutations in the edited embryos. The number of edited embryos for each genotype was also shown.

In previous studies, RNP transformation showed low off-target mutations in plants ^39,40^. One off-target sequence was identified for the crRNA targeting *CsPDS* using the CRISPR P 2.0 system ^49^. However, Sanger sequencing analyses of the 58 and 15 embryos generated by transformation of embryogenic protoplasts with ErCas12a/crRNA RNP and LbCas12aU/crRNA RNP, respectively, did not identify any mutations in the off-target homozygous sequence (Extended Data Fig. S2).

## Generation of transgene-free canker-resistant *C. sinensis* cv. Hamlin by genome editing of the canker susceptibility gene *CsLOB1*

Because LbCas12aU demonstrated superior activity in *in vitro* digestion of target sequence (Extended Data Fig. S1) and its high efficacy in genome editing of the *CsPDS* gene (Fig. 2), we used LbCas12aU/crRNA RNP in downstream studies to generate transgene-free canker-resistant *C. sinensis* cv. Hamlin by editing coding region of the canker susceptibility gene *CsLOB1*. We used one crRNA to target the 2^nd^ exon of the *CsLOB1* gene (Fig. 3A) in the RNP complex. The crRNA was carefully designed to reduce off-target homologous sites. In total 42 plantlets were regenerated (Fig. 1) and 39 lines survived in the greenhouse after micro-grafting on Carrizo. Intriguingly, Sanger sequencing analyses of the 39 regenerated lines demonstrated 38 lines were homozygous (8 lines)/biallelic (30 lines) mutants whereas only one line was wild type (Table 1), demonstrating a homozygous/biallelic mutation rate of 97.4%. The edited lines contained 12 different genotypes including 1 homozygous type (−7/-7) and 11 biallelic types (−11/-7, −11/-9, −14/-7, −4/-3, −7/-2, −7/-4, −7/-6, −7/-7 (different), −19/-7, −8/-4, −9/-6+348) (Table 1, Extended Data Figs. S3-S14). Overall, the most common mutation was 7 bp deletion (42 events), followed by −14 (10 events), −4 (7 events), −11 (5 events), −2 (4 events), −3 and −9 (2 events each), −6, −8, −19 and −6+348 (1 event each) (Table 1). The high frequency of 7 bp deletion was consistent with the genotype (−7/-7) of homozygous mutants. We further confirmed the edited lines by conducting whole genome sequencing using next generation sequencing of one representative line for each of the 12 different mutant genotypes (Table 1, Extended Data Table S1, Extended Data Figs. S3-S14). The whole genome sequencing data were in accordance with the Sanger sequencing data and confirmed the biallelic/homozygous mutations for the 12 mutant genotypes (Extended Data Figs. S3-S14). Intriguingly, one edited line contained a 348 bp insertion sequence of *C. sinensis* mitochondrial sequence at the target site of one allele of *CsLOB1*. As expected for RNP-mediated genome editing, analyses of whole genome sequencing data demonstrated that all the 12 edited lines did not contain foreign genes.

**Figure 3.**
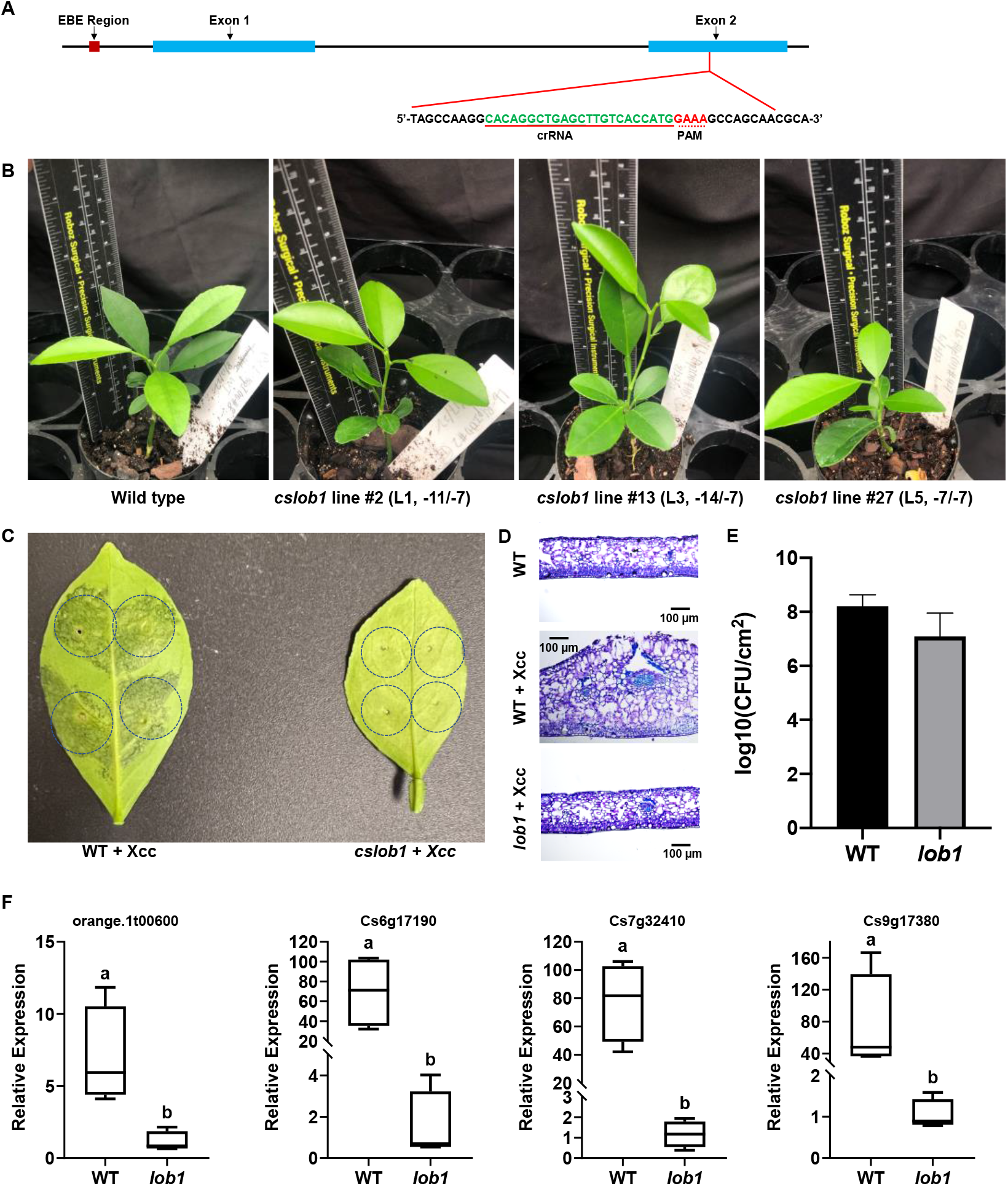
Transgene-free *cslob1* mutants of *Citrus sinensis* cv. Hamlin generated by genome editing of the *CsLOB1* gene. A. Schematic representation of the *CsLOB1* gene and crRNA. Blocks in blue indicate exons. Line fragments indicate introns. PAM: protospacer adjacent motif. B. Representative transgene-free *cslob1* mutants of *C. sinensis* cv. Hamlin grafted on Carrizo citrange *(Poncirus trifoliata* x *Citrus sinensis)*kept in greenhouse. The genotypes of the mutants were demonstrated. C. Canker symptoms on wild type *C. sinensis* cv. Hamlin and *cslob1* mutant. Fully expanded citrus leaves were inoculated with Xcc at 10^7^ CFU/ml using needleless syringes. The picture was taken at 9 days after inoculation. Six biallelic/homozygous lines were randomly selected and tested with similar results. Only one representative picture was shown. D. Thin cross-section images of C. E. Xcc titers at 9 days post-inoculation. Four biological replicates were used. The experiments were repeated at least two times with comparable results. F. Expression of *orange. 1t00600, Cs6g17190, Cs7g32410* and *Cs9g17380*, which are known to be up-regulated by CsLOB1 during Xcc infection, in the *cslob1* mutant and wild-type *C. sinensis* cv. Hamlin. *CsGAPDH*, a housekeeping gene encoding glyceraldehyde-3-phosphate dehydrogenase in citrus was used as an endogenous control. Four biological replicates were used and mean values ± SD (*n* = 4) are shown. Student’s *t* test was used for statistical analysis, different letters showed significant differences (P-value<0.05). Experiments were repeated at least two times with similar results and representative results are shown.

**Table 1.** Summary of transgene-free *CsLOB1*-edited *C. sinensis* cv. Hamlin lines generated by LbCas12aU/crRNA RNP transformation of embryogenic citrus protoplasts.

| Types of regenerated lines | Mutation types | Mutations of two alleles | Number of regenerated lines |
| --- | --- | --- | --- |
| L1 | Biallelic | -11/-7 | 4 |
| L2 | Biallelic | -7/-2 | 4 |
| L3 | Biallelic | -14/-7 | 10 |
| L4 | Biallelic | -19/-7 | 1 |
| L5 | Homozygous | -7/-7 | 8 |
| L6 | Biallelic | -7/-4 | 4 |
| L7 | Biallelic | -7/-7 (different) | 1 |
| L8 | Biallelic | -8/-4 | 1 |
| L9 | Biallelic | -7/-6 | 1 |
| L10 | Biallelic | -4/-3 | 2 |
| L11 | Biallelic | -11/-9 | 1 |
| L12 | Biallelic | -9/-6+348 | 1 |
| L13 | Wild type |  | 1 |

Next, we investigated whether our transgene-free lines contained off-target mutations at homologous sites to the targeting site. We searched for potential off-target sites of the crRNA targeting *CsLOB1* gene using the CRISPR P v2.0 program and only one homologous site that differed by up to 4 nucleotides was identified. Both amplicon deep sequencing (Extended Data Table S2) and whole genome sequencing analyses showed no off-target mutations. In addition, we further investigated whether mutations occurred in *CsLOB1* homologs. *C. sinensis* contains two *CsLOB1* functional homologs, *CsLOB2* and *CsLOB3* that share 67.9% and 71.0% identities to *CsLOB1*, respectively ^50^. Whole genome sequencing analysis showed that *CsLOB2* and *CsLOB3* sequences in the 12 edited lines were identical to their counterparts in wild type *C. sinensis* cv. Hamlin.

The biallelic/homozygous *cslob1* mutants showed no differences from wild type *C. sinensis* cv. Hamlin in growth phenotypes (Fig. 3B). As expected, Xcc infection of the biallelic/homozygous *cslob1* mutants did not cause any canker symptoms (Fig. 3C; Extended Data Fig. S15). The typical hypertrophy and hyperplasia in leaf tissues caused by Xcc were abolished by mutation of the *CsLOB1* gene (Fig. D). The canker resistance of the biallelic/homozygous *cslob1* mutants resulted from abolishing the canker symptoms rather than inhibiting Xcc growth. There were no significant differences in Xcc titers between the wild type and *cslob1* mutants (Fig. 3E; Extended Data Fig. S16). Next, we investigated the expression of *Cs7g32410* (expansin), *orange.1t00600* (3-oxo-5-alpha-steroid 4-dehydrogenase), *Cs6g17190* (RSI-1), and *Cs9g17380* (PAR1), which were known to be up-regulated by CsLOB1 during Xcc infection^50–52^, in the wild type and *cslob1* mutant. qRT-PCR analysis clearly demonstrated that expression of *orange.1t00600, Cs6g17190, Cs7g32410* and *Cs9g17380* was significantly lower in the *cslob1* mutant than in the wild type *C. sinensis* in the presence of Xcc (Fig. F), indicating mutation of *CsLOB1* abolished the induction of downstream genes of *CsLOB1* by Xcc.

## Discussion

In this study, we have generated transgene-free canker-resistant *C. sinensis* cv. Hamlin lines via Cas12a/crRNA RNP transformation of embryogenic citrus protoplasts by editing the coding region of canker susceptibility gene *CsLOB1*. Canker resistance resulting from editing the coding region of canker susceptibility *CsLOB1* is consistent with previous results in enabling canker resistance by editing the promoter region or coding regions of *LOB1* genes in grapefruit (*C. paradisi*)^27,29^, sweet orange (*C. sinensis*)^31,32^ and pummelo (*C. maxima*)^28^. Interestingly, natural variations of the effector binding elements in the *LOB1* promoter were reported to contribute to citrus canker disease resistance in *Atalantia buxifolia*, a primitive (distant-wild) citrus^53^. Mutation of the coding region or promoter region of susceptibility genes via genome editing or utilizing their natural variants have been successfully used in generating disease-resistant plants such as bacterial blight-resistant rice varieties ^54^, powdery mildew-resistant wheat ^55^, and enabling broad resistance to bacterial, oomycete, and fungal pathogens^56^.

Biotechnological approaches including transgenic expression, RNAi, and CRISPR mediated genome editing have been used in citrus genetic improvement ^13,16,17,19,20,29,57–60^. However, none of them have been adopted for commercial use despite significant improvements in different traits including resistance to diseases and shortened juvenility. The lack of success in commercialization for citrus plants generated by biotechnological approaches primarily results from their transgenic nature. Transgenic plants need to pass rigorous, lengthy, and costly regulatory approvals. The regulatory requirements by different countries/regions vary ^33,34^. In the U.S., transgenic plants are regulated by the Animal & Plant Health Inspection Service (APHIS), Environmental Protection Agency (EPA), and Food and Drug Administration (FDA). Our *CsLOB1* edited *C. sinensis* cv. Hamlin lines were generated by the RNP method that does not involve DNA fragments. Consequently, the edited *C. sinensis* lines do not contain foreign genes, which is consistent with other genome-edited plants generated by the RNP approach ^36–41^. In agreement with the low off-target efficacy of RNP-mediated genome editing ^39,40^, the *CsLOB1* edited *C. sinensis* cv. Hamlin lines do not have off-target mutations including in *CsLOB2* and *CsLOB3*, the two *CsLOB1* homologs. Importantly, the *CsLOB1* edited *C. sinensis* cv. Hamlin lines demonstrate no phenotypic differences from wild type plants except canker resistance. Mutation of the *LOB1* homolog in *Arabidopsis* was also reported to have no effect on plant phenotypes due to functional redundancy of *LOB1* gene and its homologs ^61^. Owing to the long juvenile period, we were unable to evaluate the fruit quality and yield of the edited lines, which is expected to be completed in 3 more years. Because of the aforementioned traits, the transgene-free canker-resistant *C. sinensis* cv. Hamlin lines have received regulatory approval by APHIS (Supplementary Information File 1), clearing one important hurdle for its potential use in production. In addition, enabling plant disease resistance via editing susceptibility genes is different from that via utilizing R gene. Editing of *LOB1* gene, the canker susceptibility gene, seems to have a neglectable effect on Xcc growth when inoculated using injection while abolishing the gene induction of downstream genes of *CsLOB1*, thus obliterating disease symptoms and damages. In contrast, R gene-based disease resistance usually reduces pathogen titers and causes hypersensitive response ^62^. Editing susceptibility genes such as *LOB1* to enable plant resistance probably represents a unique advantage in passing the regulation of EPA since the plants have no effects on pathogen growth, thus do not qualify as a plant-incorporated protectant with pesticidal nature which EPA regulates.

It was reported that RNP-mediated genome editing suffers from low genome editing efficacy^42^. In our study, 38 of the 39 regenerated *CsLOB1* edited lines were biallelic/homozygous mutants, demonstrating a 97.4% biallelic/homozygous mutation rate. In addition, both LbCas12aU/crRNA and ErCas12a/crRNA RNP were able to efficiently generate biallelic/homozygous *CsPDS* mutations. The high efficacy of RNP-mediated citrus genome editing suggests room for improvement and optimization for RNP genome editing in other plant species. The entire process of RNP-mediated citrus genome editing, from transformation to grafting, takes about 10 months (Fig. 1). This significantly shortens the approximately 20 years needed for citrus breeding through traditional approaches ^12^. In sum, this study develops an efficient genome editing approach for citrus using RNP, which has a significant impact on the genetic improvement of elite citrus cultivars in different traits including against some crushing challenges such as citrus Huanglongbing ^63,64^. It is anticipated that the RNP-mediated genome editing can be used in genetic improvement and genetic study of many plant species beyond citrus. In addition, this study generated the first transgene-free canker-resistant *C. sinensis* lines that are in the process of being evaluated and released to provide a sustainable and efficient solution to control citrus canker.

## Methods

### Growth conditions of citrus plants and cell culture

For *C. sinensis*, the young seedlings were grown in a greenhouse located in Citrus Research and Education Center, Lake Alfred, FL. Embryogenic callus lines of *C. sinensis* (Hamlin sweet orange) were initiated from immature ovules and maintained on Murashige and Tucker (1969, MT) medium (Phytotech Labs, Lenexa, KS, USA) ^65^ supplemented with 5.0 mg/l Kinetin (KIN) and 500 mg/l malt extract. The suspension cell culture of *C. sinensis* cv. Hamlin was maintained under dark at 22°C and sub-cultured every two weeks. The growing medium was H+H medium (MT basal medium plus 35 g/L sucrose, 0.5 g/L malt extract, 1.55 g/L glutamine, pH 5.8) ^66^. At 7-10 days after subculturing, the suspension cells were used for protoplast isolation.

### Protoplast isolation

Embryogenic *C. sinensis* cv. Hamlin protoplasts were isolated from the suspension cells after digestion with digestion solution (2.5x volume BH3 and 1.5x volume enzyme solution (0.7 M mannitol, 24 mM CaCl2, 6.15 mM MES buffer, 2.4 % (w/v) Cellulase Onozuka RS (Yakult Honsha, Minato-ku, Tokyo, Japan), 2.4 % (w/v) Macerozyme R-10 (Yakult Honsha), pH 5.6) for 16-20 hours at 28°C. After digestion, the digestion protoplast mixture was filtered with a 40 μM cell strainer (Corning, Durham, NC, USA) into a 50 ml Falcon tube, which were centrifuged at 700 rpm for 7 min. The pellets were resuspended with BH3 medium to wash the protoplast ^48^. After repeating the washing step, the protoplasts were resuspended in 2 mL BH3 medium and diluted to 1 x 10^6^ cell/mL and kept in dark at room temperature for 1 hour.

### Cas12a proteins and crRNA molecules

ErCas12a protein with a single, carboxy-terminal SV40-derived nuclear localization signal was received from Integrated DNA Technologies (IDT, Coralville, Iowa). DNA sequence encoding ErCas12a was cloned into the pET28a vector by Gibson assembly. For protein expression, a single transformed *E. coli* BL21(DE3) colony was inoculated into LB medium supplemented with Kanamycin (25 μg/mL), and grown overnight at 37°C, 250 rpm. The overnight culture was transferred to terrific broth medium containing 0.5% glucose and 25 μg/mL kanamycin, grown at 37 °C, 250 rpm for ~2–3 h until OD_600_ reached 0.6. The culture was chilled at 4 °C for 30 min prior to induction with 1 mM IPTG, and further incubated at 18 °C, 250 rpm for 12~18 h. The recombinant ErCas12a protein was purified as previously described ^67^. Briefly, *E. coli* cells were harvested by centrifugation, and homogenized with Emulsiflex-C3 high-pressure homogenizer (Avestin, Ottawa ON, Canada). The ErCas12a protein in clarified lysate was sequentially purified using immobilized metal affinity chromatography (HisTrap HP, GE Healthcare) and heparin chromatography (HiTrap Heparin HP, GE Healthcare). Purified protein was concentrated and dialyzed against storage buffer (20 mM TrisHCl, 300 mM NaCl, 0.1 mM EDTA, 50% Glycerol, 1 mM DTT, pH 7.4) overnight. The protein concentration was measured by NanoDrop using extinction coefficient at 143,940 M^-1^cm^-1^, diluted to 60 μM, and stored at −20 °C. Alt-RL.b. Cas12a (Cpf1) Ultra (LbCas12aU) protein with a single carboxy-terminal SV40-derived nuclear localization signal was purchased from IDT. crRNAs targeting *CsPDS* or *CsLOB1* genes were selected by manually searching for the PAM site “TTTV”. crRNAs (Extended Data Table S2) targeting *CsPDS* or *CsLOB1* genes were synthesized by IDT and diluted to 0.05 nmol/μl by RNase-free water.

### *in vitro* digestion

ErCas12a (1 μg) or LbCas12aU (1 μg) protein and 1 μg crRNA were assembled in 1X Nuclease Reaction Buffer (New England BioLabs, Ipswich, MA, USA) at room temperature for 10 minutes. Then 100 ng DNA fragments were added to the mixture in a total volume of 30 μL. Digested DNA products were run using 2% agarose gel after 30 min digestion at 37°C.

### Transformation of embryogenic citrus protoplast with RNP using the PEG method and plant regeneration

For RNP assembly, 0.27 nmol ErCas12a/LbCas12aU protein and 0.45 nmol crRNA were assembled in 1 X Nuclease Reaction Buffer (NEB). The protein and RNA were mixed and incubated for 10 minutes at room temperature and used for transformation of embryogenic *C. sinensis* cv. Hamlin protoplasts using the PEG method ^48^.

For each transfection reaction, 1 ml protoplast cells, 20 μL preassembled RNP, and 1 ml PEG-CaCl_2_ (0.4 M mannitol, 100 mM CaCl_2_, and 40% PEG-4000) were mixed and kept at room temperature for 15 min in dark followed by washing with BH3 medium twice. The RNP-transformed embryogenic citrus protoplasts were used for plant regeneration (Fig. 1).

### Mutation detection based on PCR amplification, and Sanger sequencing

Genomic DNA was extracted from leaves of wild type or *cslob1* mutants of *C. sinensis* cv. Hamlin. For *cspds* mutants, genomic DNA was extracted from embryos. Primers used for PCR were listed in the Extended Data Table S2. CloneAmp HiFi PCR Premix (Takara Bio USA, San Jose, CA, USA) was used for PCR amplification following the manufacturer’s instructions using the following protocol: 98°C for 30 s; followed by 40 cycles at 98°C for 10 s, 54°C for 10 s, and 72°C for 45 s; followed by a final extension at 72°C for 5 min. PCR amplicons were sequenced directly using the amplifying primers or cloned with Zero Blunt TOPO PCR Cloning Kit (Thermo Fisher, San Jose, CA, USA) and transformed into Stellar Competent Cells (Takara). M13-F (GTAAAACGACGGCCAGTG) and M13-R (CAGGAAACAGCTATGACC) were used for single colony PCR amplification and Sanger sequencing.

### DNA Library construction, sequencing, and data analysis

Following the manufacturer’s protocol of short read DNA sequencing from Illumina ^68^, the library was prepared. After quality control, quantification, and normalization of the DNA libraries, 150 bp paired-end reads were generated using the Illumina NovaSeq 6000 platform according to the manufacturer’s instructions at Novogene. The raw paired-end reads were filtered to remove low quality reads using fastp program version 0.22.0 ^69^. On average, more than 21.45 Gb of high-quality data was generated for each edited sweet orange plant sample (Extended Data Table S1). To identify the mutations (single nucleotide polymorphisms, deletions and insertions) for the mutated plant genomes, the high-quality paired-end short genomic reads were mapped to sweet orange (*C. sinensis*) ^70^ reference genome using Bowtie2 software version 2.2.6 ^71^. Based on the mapping results, mutations were detected using the SAMtools package version 1.2 ^72^ and deepvariant program version 1.4.0 ^73^. The generated mutations were filtered by quality and sequence depth (mapping quality > 10 and mapping depth >10). The mutations of target site were visualized using the Integrative Genomics Viewer (IGV) software version 2.15.4 ^74^. The high-quality paired-end short reads were further used to detect foreign DNA sequences. The off-target sites were predicted by using CRISPR-P 2.0 program ^49^ and aligning target sequence with whole genome using blast program. Based on the mapping results, mutations of off-target sites were detected using the SAMtools package version 1.2 and deepvariant program version 1.4.0.

### Quantitative reverse-transcription PCR (qRT-PCR)

Xcc strain 306 was infiltrated into wild type *C. sinensis* cv. Hamlin and transgene-free *cslob1* mutants at the concentration of 1×10^7^ cfu/mL. The infiltration-area of the leaf samples were collected at 9 days post-inoculation (dpi) for RNA isolation. Four biological repeats were used with one leaf as one biological replicate. Total RNA was extracted by TRIzo1 Reagent (Thermo-Fisher) following the manufacturer’s instructions. cDNA was synthesized by qScript cDNA SuperMix (Quantabio, Beverly, MA, USA). Primers used for qRT-PCR were listed in Extended Data Table S2. Briefly, the real-time PCR was performed with QuantiStudio3 (Thermo-Fisher) using SYBR Green Real-Time PCR Master Mixes (Thermo-Fisher) in a 10 μL reaction. The standard amplification protocol was 95°C for 3 min followed by 40 cycles of 95°C 15 s, 60°C for 60 s. *CsGAPDH* was used as an endogenous control. All reactions were performed in triplicate. Relative gene expression and statistical analysis were calculated using the 2^-ΔΔCT^method ^75^. qRT-PCR was repeated twice with similar results.

### Microscopy analysis

The infiltration areas of Xcc-infiltrated wild type *C. sinensis* cv. Hamlin and *cslob1* mutant leaves and non-inoculated wild type Hamlin leaves were cut with sterilized blades and fixed in 4% paraformaldehyde for at least 2 hours. The specimen was dehydrated and embedded in paraffin chips. The paraffin chips were sectioned with Leica 2155 microtome and the thickness of the cut ribbon was 8 μm. The ribbons were located on the glass slides and incubated at 37°C overnight to be heat fixed. Followed by the dewaxing and rehydrating process, the slides were stained with 0.05% Toluidine blue for 30 seconds, then rinsed in ddH2O, dehydrated, and added one drop of mounting media, covered with a coverslip. After solidifying for one hour, the photos of the slides were taken with Leica LasX software (Leica Biosystems Inc., Lincolnshire, IL, USA) under the bright-field microscope (Olympus BX61; Olympus Corporation, Shinjuku City, Tokyo, Japan).

### Xcc growth assay

Leaf discs (0.5 cm in diameter) punched from the inoculated plant leaves were ground in 0.2 mL sterilized H2O. 100 μL serial dilutions of the grinding suspensions were spread on NA plates (dilutions ranging from 10^-1^ to 10^-6^). Bacterial colonies were counted after 48 h and the number of CFU per mL suspension solution of leaf disc was calculated and presented with Prism GraphPad software.

## Data availability

The raw reads of genome resequencing for sweet orange plants were deposited in the NCBI Bioproject database under the accession number PRJNA931574. The reference genome of sweet orange was downloaded from public citrus genome database CPBD: Citrus Pan-genome to Breeding Database (http://citrus.hzau.edu.cn/index.php).

## Acknowledgments

We thank Wang lab members for constructive suggestions and insightful discussions. This project was supported by funding from Florida Citrus Initiative Program, Citrus Research and Development Foundation, U.S. Department of Agriculture National Institute of Food and Agriculture grants 2022-70029-38471, 2021-67013-34588, 2018-70016-27412 and 2016-70016-24833, FDACS Specialty Crop Block Grant Program (N. Wang).

## Author Contributions

N. W. conceptualized, designed the experiments and supervised the project. H. S. and Y. C. W. performed the experiments. A. O. and J. G. provided citrus suspension culture, L. Y. Z. and C. V. provided Cas12a proteins. J.X., and H.S. performed bioinformatics and statistical analyses. N. W., H. S., J. X., and Y. C. W. wrote the manuscript with input from all co-authors.

## Competing interests

N. W., H. S. and Y. C. W. filed a PCT patent application based on the results reported in this paper. All other authors declare no competing financial interests.

**Correspondence and requests** for materials should be addressed to N. Wang.

## Figure legends

**Extended Data Fig. S1.** Evaluate the crRNA guided endonuclease activity of Cas12a *in vitro*. A. Schematic representation of the *CsPDS* gene (orange1.1t02361) and crRNA. Blocks in blue indicate exons. Line fragments indicate introns. PAM: protospacer adjacent motif. B & C. *in vitro* digestion of DNA fragments using ErCas12a (B) and LbCAs12aU (C). A DNA fragment (555 bp) of the *CsPDS* gene containing the crRNA target site as depicted in A was digested with ErCas12a (B) or LbCas12aU (C). After 30 min, DNA electrophoresis was run using 2% agarose gel.

**Extended Data Fig. S2.** Off-target analysis by PCR amplification, cloning and Sanger sequencing of embryos generated after ErCas12a and LbCas12aU RNP transformation. 58 embryos generated after ErCas12a RNP transformation of embryogenic protoplasts and 15 embryos generated after LbCas12aU RNA transformation of embryogenic protoplasts were tested. The sequencing result of representative embryos from each genotype were shown. No off-target activity was detected in these embryos.

**Extended Data Fig. S3.** Sequencing confirmation of the transgene-free *CsLOB1*-edited *C. sinensis* cv. Hamlin line L1 generated by LbCas12aU/crRNA RNP transformation of embryogenic citrus protoplasts. A. Sequencing confirmation of *CsLOB1* based on PCR amplification and cloning. The representative chromatograms of *CsLOB1* edited *C. sinensis* cv. Hamlin lines. The mutations of both alleles of *CsLOB1* were shown for each line. x indicates number of colonies sequenced. Nucleotide in red indicates crRNA. The underlined GAAA indicates protospacer-adjacent motif (PAM). -: deletion. +: insertion. B. Whole genome sequencing of the edited lines using next generation sequencing. The bases of target site were highlighted by colors other than black. There were two types of deletions of target site, including type I (−11 bp deletion) and Type II (−7 bp deletion), which were shown by horizontal bar chart. The vertical bar chart showed the sequence depth for each nucleotide. **Extended Data Fig. S4.** Sequencing confirmation of the transgene-free *CsLOB1-edited C. sinensis* cv. Hamlin line L2 generated by LbCas12aU/crRNA RNP transformation of embryogenic citrus protoplasts. A. Sequencing confirmation of *CsLOB1* based on PCR amplification and cloning. The representative chromatograms of *CsLOB1* edited *C. sinensis* cv. Hamlin lines. The mutations of both alleles of *CsLOB1* were shown for each line. x indicates number of colonies sequenced. Nucleotide in red indicates crRNA. The underlined GAAA indicates protospacer-adjacent motif (PAM). -: deletion. +: insertion. B. Whole genome sequencing of the edited lines using next generation sequencing. The bases of target site were highlighted by colors other than black. There were two types of deletions of target site, including type I (−7 bp deletion) and Type II (−2 bp deletion), which were shown by horizontal bar chart. The vertical bar chart showed the sequence depth for each nucleotide.

**Extended Data Fig. S5.** Sequencing confirmation of the transgene-free *CsLOB1*-edited *C. sinensis* cv. Hamlin line L3 generated by LbCas12aU/crRNA RNP transformation of embryogenic citrus protoplasts. A. Sequencing confirmation of *CsLOB1* based on PCR amplification and cloning. The representative chromatograms of *CsLOB1* edited *C. sinensis* cv. Hamlin lines. The mutations of both alleles of *CsLOB1* were shown for each line. x indicates number of colonies sequenced. Nucleotide in red indicates crRNA. The underlined GAAA indicates protospacer-adjacent motif (PAM). -: deletion. +: insertion. B. Whole genome sequencing of the edited lines using next generation sequencing. The bases of target site were highlighted by colors other than black. There were two types of deletions of target site, including type I (−14 bp deletion) and Type II (−7 bp deletion), which were shown by horizontal bar chart. The vertical bar chart showed the sequence depth for each nucleotide.

**Extended Data Fig. S6.** Sequencing confirmation of the transgene-free *CsLOB1*-edited *C. sinensis* cv. Hamlin line L4 generated by LbCas12aU/crRNA RNP transformation of embryogenic citrus protoplasts. A. Sequencing confirmation of *CsLOB1* based on PCR amplification and cloning. The representative chromatograms of *CsLOB1* edited *C. sinensis* cv. Hamlin lines. The mutations of both alleles of *CsLOB1* were shown for each line. x indicates number of colonies sequenced. Nucleotide in red indicates crRNA. The underlined GAAA indicates protospacer-adjacent motif (PAM). -: deletion. +: insertion. B. Whole genome sequencing of the edited lines using next generation sequencing. The bases of target site were highlighted by colors other than black. There were two types of deletions of target site, including type I (−7 bp deletion) and Type II (−19 bp deletion), which were shown by horizontal bar chart. The vertical bar chart showed the sequence depth for each nucleotide.

**Extended Data Fig. S7.** Sequencing confirmation of the transgene-free *CsLOB1*-edited *C. sinensis* cv. Hamlin line L5 generated by LbCas12aU/crRNA RNP transformation of embryogenic citrus protoplasts. A. Sequencing confirmation of *CsLOB1* based on PCR amplification and cloning. The representative chromatograms of *CsLOB1* edited *C. sinensis* cv. Hamlin lines. The mutations of both alleles of *CsLOB1* were shown for each line. x indicates number of colonies sequenced. Nucleotide in red indicates crRNA. The underlined GAAA indicates protospacer-adjacent motif (PAM). -: deletion. +: insertion. B. Whole genome sequencing of the edited lines using next generation sequencing. The bases of target site were highlighted by colors other than black. There was only one type of deletion of target site, 7 bp deletion, which was shown by the horizontal bar chart. The vertical bar chart showed the sequence depth for each nucleotide.

**Extended Data Fig. S8.** Sequencing confirmation of the transgene-free *CsLOB1*-edited *C. sinensis* cv. Hamlin line L6 generated by LbCas12aU/crRNA RNP transformation of embryogenic citrus protoplasts. A. Sequencing confirmation of *CsLOB1* based on PCR amplification and cloning. The representative chromatograms of *CsLOB1* edited *C. sinensis* cv. Hamlin lines. The mutations of both alleles of *CsLOB1* were shown for each line. x indicates number of colonies sequenced. Nucleotide in red indicates crRNA. The underlined GAAA indicates protospacer-adjacent motif (PAM). -: deletion. +: insertion. B. Whole genome sequencing of the edited lines using next generation sequencing. The bases of target site were highlighted by colors other than black. There were two types of deletions of target site, including type I (−7 bp deletion) and Type II (−4 bp deletion), which were shown by the horizontal bar chart. The vertical bar chart showed the sequence depth for each nucleotide.

**Extended Data Fig. S9**. Sequencing confirmation of the transgene-free *CsLOB1-edited C. sinensis* cv. Hamlin line L7 generated by LbCas12aU/crRNA RNP transformation of embryogenic citrus protoplasts. A. Sequencing confirmation of *CsLOB1* based on PCR amplification and cloning. The representative chromatograms of *CsLOB1* edited *C. sinensis* cv. Hamlin lines. The mutations of both alleles of *CsLOB1* were shown for each line. x indicates number of colonies sequenced. Nucleotide in red indicates crRNA. The underlined GAAA indicates protospacer-adjacent motif (PAM). -: deletion. +: insertion. B. Whole genome sequencing of the edited lines using next generation sequencing. The bases of target site were highlighted by colors other than black. There were two types of 7 bp deletions of target site which were shown by the horizontal bar chart. The vertical bar chart showed the sequence depth for each nucleotide.

**Extended Data Fig. S10.** Sequencing confirmation of the transgene-free *CsLOB1*-edited *C. sinensis* cv. Hamlin line L8 generated by LbCas12aU/crRNA RNP transformation of embryogenic citrus protoplasts. A. Sequencing confirmation of *CsLOB1* based on PCR amplification and cloning. The representative chromatograms of *CsLOB1* edited *C. sinensis* cv. Hamlin lines. The mutations of both alleles of *CsLOB1* were shown for each line. x indicates number of colonies sequenced. Nucleotide in red indicates crRNA. The underlined GAAA indicates protospacer-adjacent motif (PAM). -: deletion. +: insertion. B. Whole genome sequencing of the edited lines using next generation sequencing. The bases of target site were highlighted by colors other than black. There were two types of deletions of target site, including type I (−8 bp deletion) and Type II (−4 bp deletion), which were shown by the horizontal bar chart. The vertical bar chart showed the sequence depth for each nucleotide.

**Extended Data Fig. S11.** Sequencing confirmation of the transgene-free *CsLOB1*-edited *C. sinensis* cv. Hamlin line L9 generated by LbCas12aU/crRNA RNP transformation of embryogenic citrus protoplasts. A. Sequencing confirmation of *CsLOB1* based on PCR amplification and cloning. The representative chromatograms of *CsLOB1* edited *C. sinensis* cv. Hamlin lines. The mutations of both alleles of *CsLOB1* were shown for each line. x indicates number of colonies sequenced. Nucleotide in red indicates crRNA. The underlined GAAA indicates protospacer-adjacent motif (PAM). -: deletion. +: insertion. B. Whole genome sequencing of the edited lines using next generation sequencing. The bases of target site were highlighted by colors other than black. There were two types of deletions of target site, including type I (−7 bp deletion) and Type II (−6 bp deletion), which were shown by the horizontal bar chart. The vertical bar chart showed the sequence depth for each nucleotide.

**Extended Data Fig. S12.** Sequencing confirmation of the transgene-free *CsLOB1*-edited *C. sinensis* cv. Hamlin line L10 generated by LbCas12aU/crRNA RNP transformation of embryogenic citrus protoplasts. A. Sequencing confirmation of *CsLOB1* based on PCR amplification and cloning. The representative chromatograms of *CsLOB1* edited *C. sinensis* cv. Hamlin lines. The mutations of both alleles of *CsLOB1* were shown for each line. x indicates number of colonies sequenced. Nucleotide in red indicates crRNA. The underlined GAAA indicates protospacer-adjacent motif (PAM). -: deletion. +: insertion. B. Whole genome sequencing of the edited lines using next generation sequencing. The bases of target site were highlighted by colors other than black. There were two types of deletions of target site, including type I (−3 bp deletion) and Type II (−4 bp deletion), which were shown by the horizontal bar chart. The vertical bar chart showed the sequence depth for each nucleotide.

**Extended Data Fig. S13.** Sequencing confirmation of the transgene-free *CsLOB1*-edited *C. sinensis* cv. Hamlin line L11 generated by LbCas12aU/crRNA RNP transformation of embryogenic citrus protoplasts. A. Sequencing confirmation of *CsLOB1* based on PCR amplification and cloning. The representative chromatograms of *CsLOB1* edited *C. sinensis* cv. Hamlin lines. The mutations of both alleles of *CsLOB1* were shown for each line. x indicates number of colonies sequenced. Nucleotide in red indicates crRNA. The underlined GAAA indicates protospacer-adjacent motif (PAM). -: deletion. +: insertion. B. Whole genome sequencing of the edited lines using next generation sequencing. The bases of target site were highlighted by colors other than black. There were two types of deletions of target site, including type I (−9 bp deletion) and Type II (−11 bp deletion), which were shown by the horizontal bar chart. The vertical bar chart showed the sequence depth for each nucleotide.

**Extended Data Fig. S14.** Sequencing confirmation of the transgene-free *CsLOB1*-edited *C. sinensis* cv. Hamlin line L12 generated by LbCas12aU/crRNA RNP transformation of embryogenic citrus protoplasts. Sequencing confirmation of *CsLOB1* based on PCR amplification and cloning. The representative chromatograms of *CsLOB1* edited *C. sinensis* cv. Hamlin lines. The mutations of both alleles of *CsLOB1* were shown for each line. x indicates number of colonies sequenced. Nucleotide in red indicates crRNA. The underlined GAAA indicates protospacer-adjacent motif (PAM). -: deletion. +: insertion. For the L12 genotype, one allele contained both 6 bp deletion and 348 bp insertion of *C. sinensis* mitochondrial sequence. Only part of the insertion was shown. B. One allele of *CsLOB1* sequence in L12 showing insertion sequence and 6 bp deletion (CACAGG).

**Extended Data Fig. S15.** Canker symptoms on wild type *C. sinensis* cv. Hamlin and *cslob1*mutants. Fully expanded citrus leaves were inoculated with Xcc at 10^7^ CFU/ml using needleless syringes. The picture was taken at 9 days after inoculation.

**Extended Data Fig. S16.** *Xanthomonas citri* subsp. citri growth in wild type and transgene-free *cslob1* mutant of *Citrus sinensis* cv. Hamlin. Fully expanded citrus leaves were inoculated with Xcc at 10^7^ CFU/ml using needleless syringes. Three biological replicates were used. DPI: Days post-inoculation. 0 DPI indicates right after inoculation.

**Extended Data Table S1.** Summary of transgene-free *CsLOB1*-edited *C. sinensis* cv. Hamlin lines generated by LbCas12aU/crRNA RNP transformation of embryogenic citrus protoplasts

**Extended Data Table S2.** Primers, and crRNAs sequences

**Supplementary Information File 1**. Regulatory approval of transgene-free canker-resistant *C. sinensis* cv. Hamlin lines by APHIS

