## Supplementary Information for "Efficient generation of transgene-free canker-resistant *Citrus sinensis* using Cas12a/crRNA ribonucleoprotein"

**Extended Data Table S1.** Summary of genomic sequencing of *CsLOB1*-edited *C. sinensis* cv. Hamlin lines generated by LbCas12aU RNP transformation of embryogenic citrus protoplasts

| Types of regenerated lines | Raw data (Gb) | High quality data (Gb) |
|---|---|---|
| L1 | 23.29 | 22.79 |
| L2 | 22.56 | 22.35 |
| L3 | 21.74 | 21.22 |
| L4 | 17.23 | 16.91 |
| L5 | 20.53 | 20.15 |
| L6 | 23.50 | 23.07 |
| L7 | 23.61 | 23.12 |
| L8 | 27.81 | 27.14 |
| L9 | 19.15 | 18.76 |
| L10 | 18.53 | 18.24 |
| L11 | 21.79 | 21.34 |
| L12 | 21.51 | 21.09 |
| Wild type | 23.07 | 22.61 |

**Extended Data Table S2.** Primers, and crRNAs sequences

| Name | Sequences (5’------3’) |
|---|---|
| F-LOB1-offtarget | TCAGCATGCGTTTTTGCTGT |
| R-LOB1-offtarget | CTTGATCTCCTCTACTTTGGGG |
| F-PDS-offtarget | GCATATAGCTCGCTCGGCAA |
| R-PDS-offtarget | CGTTGTCGTCAATGATTAACTCG |
| Primers for qRT-PCR | |
| F-Cs7g32410 | GCCTCAGGAACAATGGGAGG |
| R-Cs7g32410 | CCGTGTTAACGCCGTATCCT |
| F-Cs6g17190 | CTCTCGCAGCTCCATTCTGT |
| R-Cs6g17190 | GTCGCCGAACACCGATAAGA |
| F-1t00600 | CTGGCGCTTCAACGATATGC |
| R-1t00600 | GTGAGAGGTAGACGGCGAAG |
| F-Cs9g17380 | CTTTGCAGTGGTGGCTCTTG |
| R-Cs9g17380 | TTTGGTCAAGGCTCTCGCAT |
| F1-RT-LOB1 | CCACCAACCGAACCATACAA |
| R1-RT-LOB1 | CCATGCTGCTCACTGCATCT |
| F2-RT-LOB1 | AAGGCACAGGCTGAGCTTGT |
| R2-RT-LOB1 | AAGACTTGTTCTTGAGATTGTGCCA |
| F3-RT-LOB1 | GGCACAATCTCAAGAACAAGTCT |
| R3-RT-LOB1 | GGCTCCCAAGCTGATCCAAT |

| | |
|---|---|
| F-CsGAPDH | GGAAGGTCAAGATCGGAATCAA |
| R-CsGAPDH | CGTCCCTCTGCAAGATGACTCT |
| crRNA sequences | |
| PDS (BccI) | CGAGATAGTGAACCGATGGGTCA |
| LOB1-3(BlpI) | CATGGTGACAAGCTCAGCCTGTG |

Note: The restriction enzyme digestion sites were included in the parenthesis

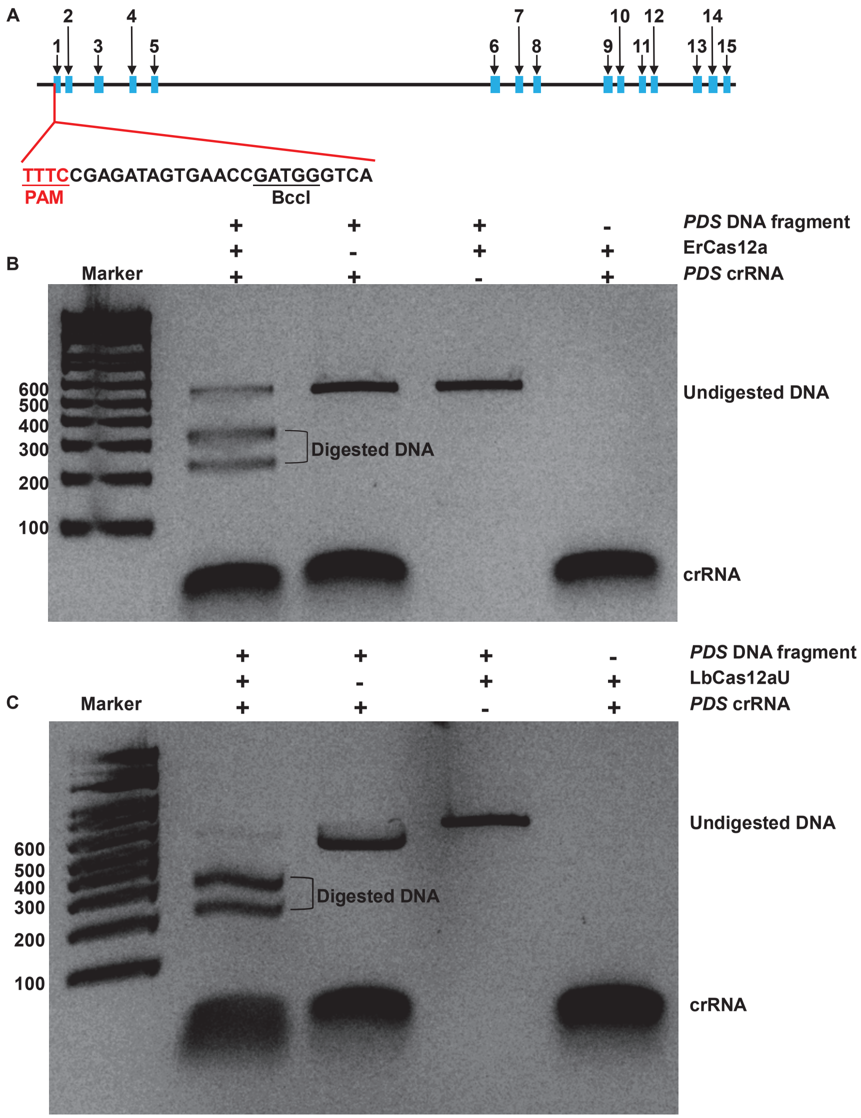

**Extended Data Fig. S1.** Test the crRNA guided endonuclease activity of Cas12a *in vitro*. A. Schematic representation of the *CsPDS* gene (orange1.1t02361) and crRNA. Blocks in blue indicate exons. Line fragments indicate introns. PAM: protospacer adjacent motif. B & C. *In vitro* digestion of DNA fragments using ErCas12a (B) and LbCAs12aU (C). A DNA fragment (555 bp) of the *CsPDS* gene containing the crRNA target site as depicted in A was digested with ErCas12a (B) or LbCas12aU. After 30 min, DNA electrophoresis was run using 2% agarose gel.

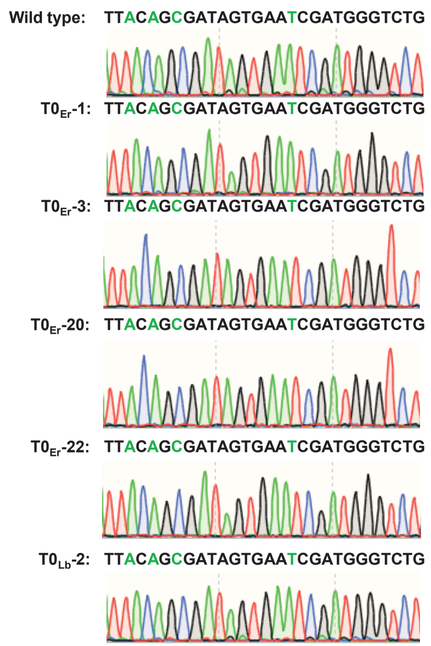

**Extended Data Fig. S2.** Off-target analysis by PCR amplification, cloning and sequencing of embryos generated after ErCas12a and LbCas12aU RNP transformation. 58 embryos generated after ErCas12a RNP transformation of embryogenic protoplasts and 15 embryos generated after LbCas12aU RNA transformation of embryogenic protoplasts were tested. The sequencing result of representative embryos from each genotype were shown. No off-target activity was detected in these embryos. 12 colonies were sequenced for each line with same results.

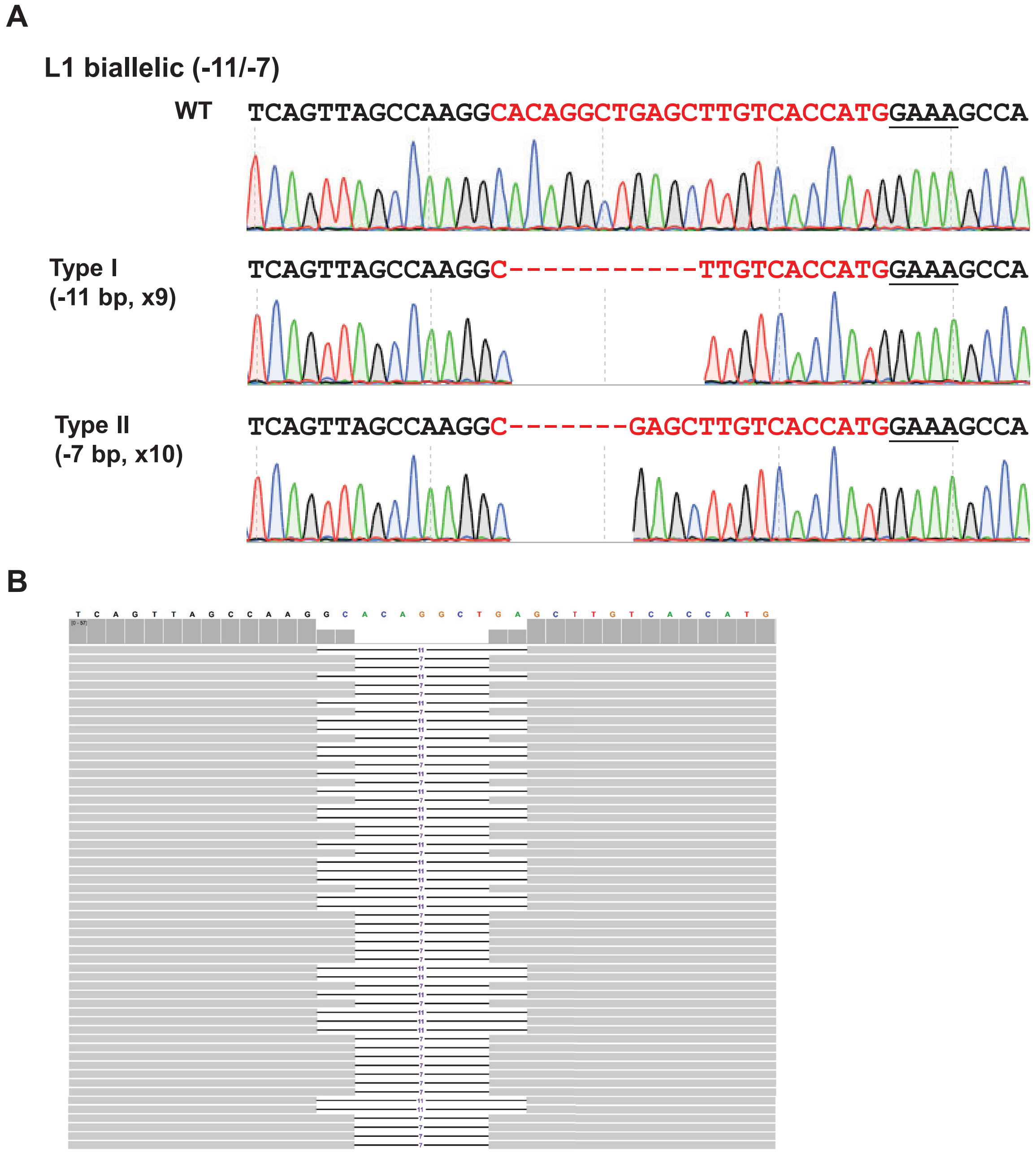

**Extended Data Fig. S3.** Sequencing confirmation of the transgene-free *CsLOB1*-edited *C. sinensis* cv. Hamlin line L1 generated by LbCas12aU/crRNA RNP transformation of embryogenic citrus protoplasts. A. Sequencing confirmation of *CsLOB1* based on PCR amplification and cloning. The representative chromatograms of *CsLOB1* edited *C. sinensis* cv. Hamlin lines. The mutations of both alleles of *CsLOB1* were shown for each line. x indicates number of colonies sequenced. Nucleotide in red indicates crRNA. The underlined GAAA indicates protospacer-adjacent motif (PAM). -: deletion. +: insertion. B. Whole genome sequencing of the edited lines using next generation sequencing. The bases of target site were highlighted by colors other than black. There were two types of deletions of target site, including type I (-11 bp deletion) and Type II (-7 bp deletion), which were shown by horizontal bar chart. The vertical bar chart showed the sequence depth for each nucleotide.

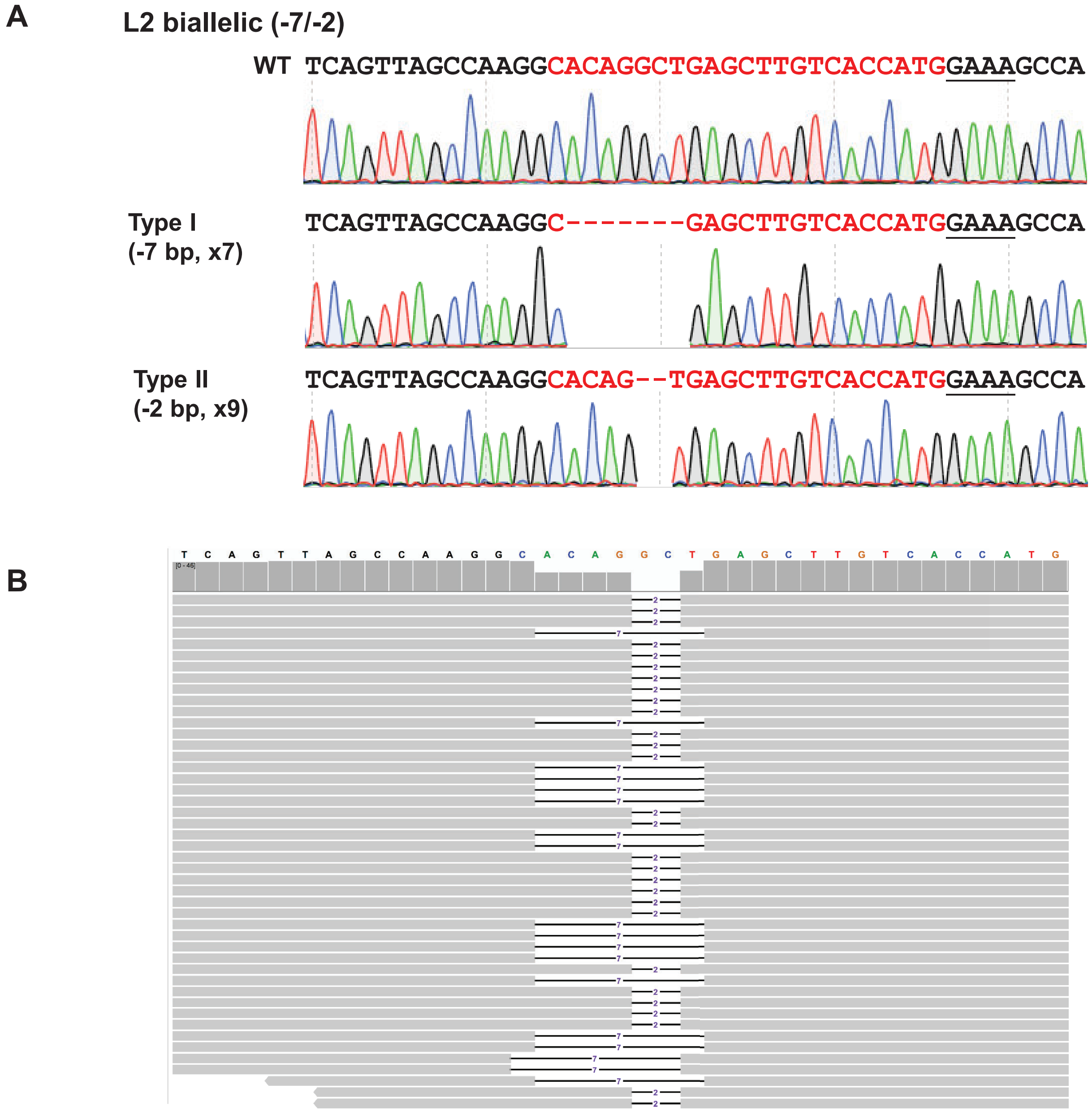

**Extended Data Fig. S4.** Sequencing confirmation of the transgene-free *CsLOB1*-edited *C. sinensis* cv. Hamlin line L2 generated by LbCas12aU/crRNA RNP transformation of embryogenic citrus protoplasts. A. Sequencing confirmation of *CsLOB1* based on PCR amplification and cloning. The representative chromatograms of *CsLOB1* edited *C. sinensis* cv. Hamlin lines. The mutations of both alleles of *CsLOB1* were shown for each line. x indicates number of colonies sequenced. Nucleotide in red indicates crRNA. The underlined GAAA indicates protospacer-adjacent motif (PAM). -: deletion. +: insertion. B. Whole genome sequencing of the edited lines using next generation sequencing. The bases of target site were highlighted by colors other than black. There were two types of deletions of target site, including type I (-7 bp deletion) and Type II (-2 bp deletion), which were shown by horizontal bar chart. The vertical bar chart showed the sequence depth for each nucleotide.

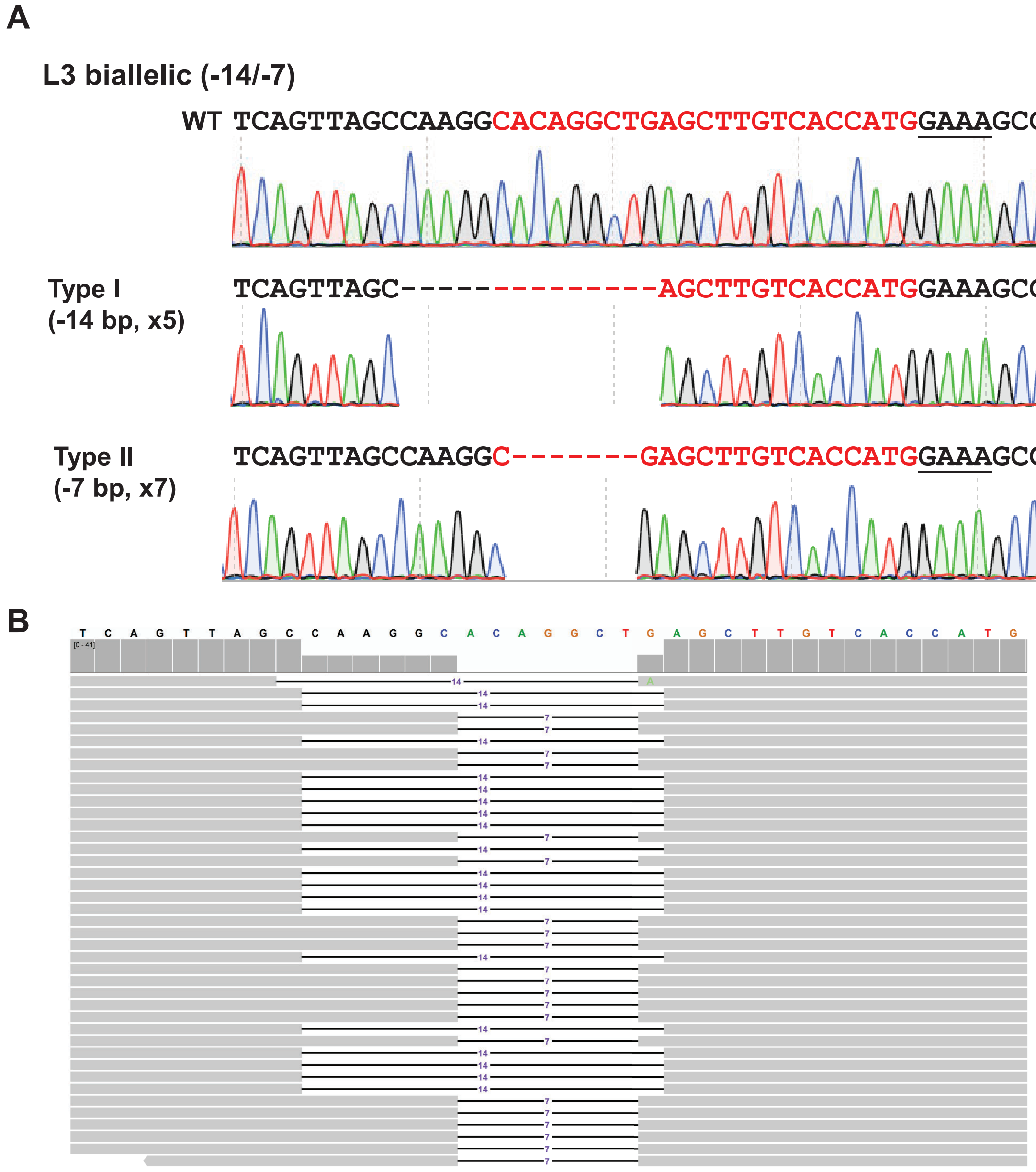

**Extended Data Fig. S5.** Sequencing confirmation of the transgene-free *CsLOB1*-edited *C. sinensis* cv. Hamlin line L3 generated by LbCas12aU/crRNA RNP transformation of embryogenic citrus protoplasts. A. Sequencing confirmation of *CsLOB1* based on PCR amplification and cloning. The representative chromatograms of *CsLOB1* edited *C. sinensis* cv. Hamlin lines. The mutations of both alleles of *CsLOB1* were shown for each line. x indicates number of colonies sequenced. Nucleotide in red indicates crRNA. The underlined GAAA indicates protospacer-adjacent motif (PAM). -: deletion. +: insertion. B. Whole genome sequencing of the edited lines using next generation sequencing. The bases of target site were highlighted by colors other than black. There were two types of deletions of target site, including type I (-14 bp deletion) and Type II (-7 bp deletion), which were shown by horizontal bar chart. The vertical bar chart showed the sequence depth for each nucleotide.

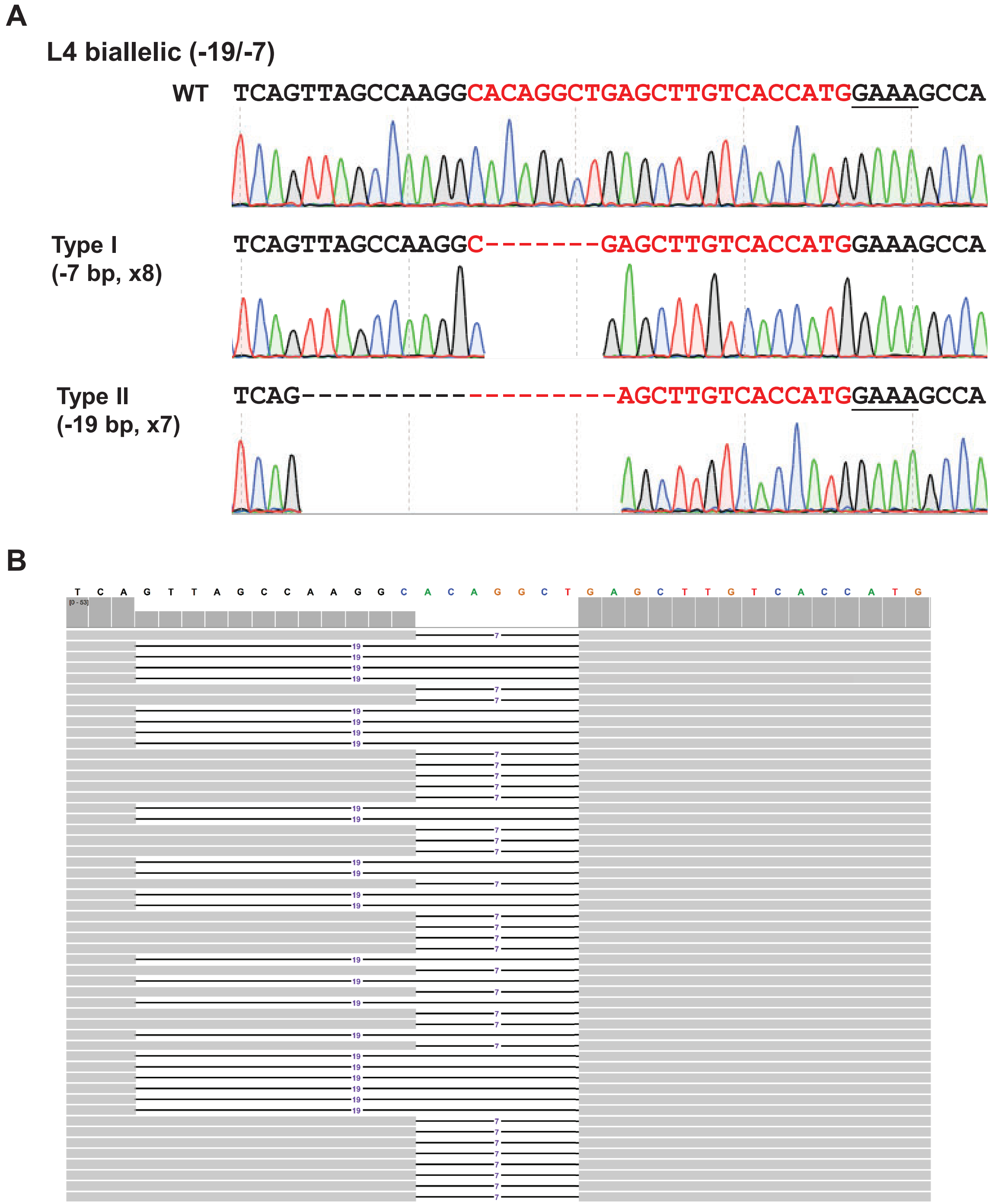

**Extended Data Fig. S6.** Sequencing confirmation of the transgene-free *CsLOB1*-edited *C. sinensis* cv. Hamlin line L4 generated by LbCas12aU RNP transformation of embryogenic citrus protoplasts. A. Sequencing confirmation of *CsLOB1* based on PCR amplification and cloning. The representative chromatograms of *CsLOB1* edited *C. sinensis* cv. Hamlin lines. The mutations of both alleles of *CsLOB1* were shown for each line. x indicates number of colonies sequenced. Nucleotide in red indicates crRNA. The underlined GAAA indicates protospacer-adjacent motif (PAM). -: deletion. +: insertion. B. Whole genome sequencing of the edited lines using next generation sequencing. The bases of target site were highlighted by colors other than black. There were two types of deletions of target site, including type I (-7 bp deletion) and Type II (-19 bp deletion), which were shown by horizontal bar chart. The vertical bar chart showed the sequence depth for each nucleotide.

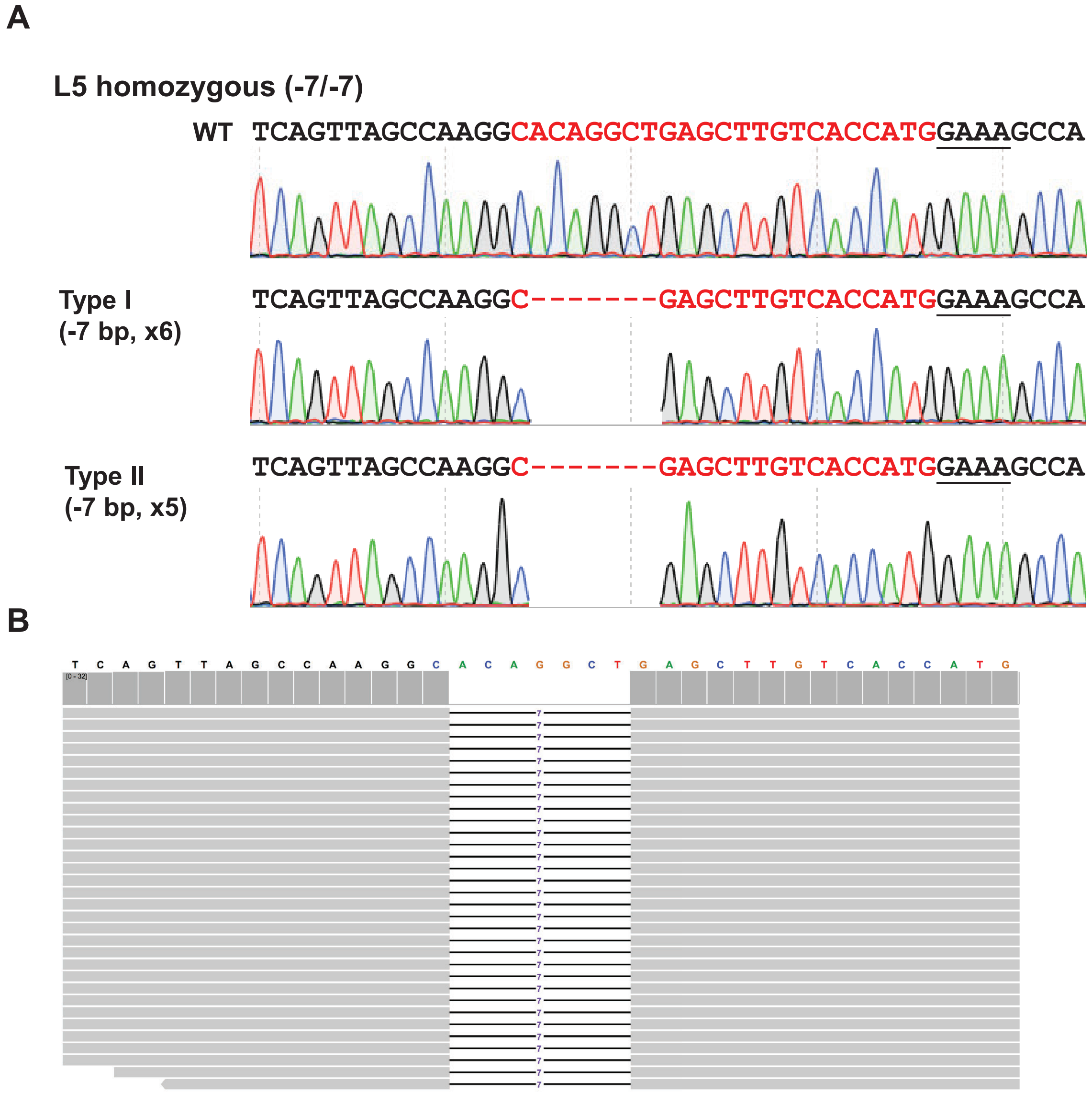

**Extended Data Fig. S7.** Sequencing confirmation of the transgene-free *CsLOB1*-edited *C. sinensis* cv. Hamlin line L5 generated by LbCas12aU/crRNA RNP transformation of embryogenic citrus protoplasts. A. Sequencing confirmation of *CsLOB1* based on PCR amplification and cloning. The representative chromatograms of *CsLOB1* edited *C. sinensis* cv. Hamlin lines. The mutations of both alleles of *CsLOB1* were shown for each line. x indicates number of colonies sequenced. Nucleotide in red indicates crRNA. The underlined GAAA indicates protospacer-adjacent motif (PAM). -: deletion. +: insertion. B. Whole genome sequencing of the edited lines using next generation sequencing. The bases of target site were highlighted by colors other than black. There was only one type of deletion of target site, 7 bp deletion, which was shown by the horizontal bar chart. The vertical bar chart showed the sequence depth for each nucleotide.

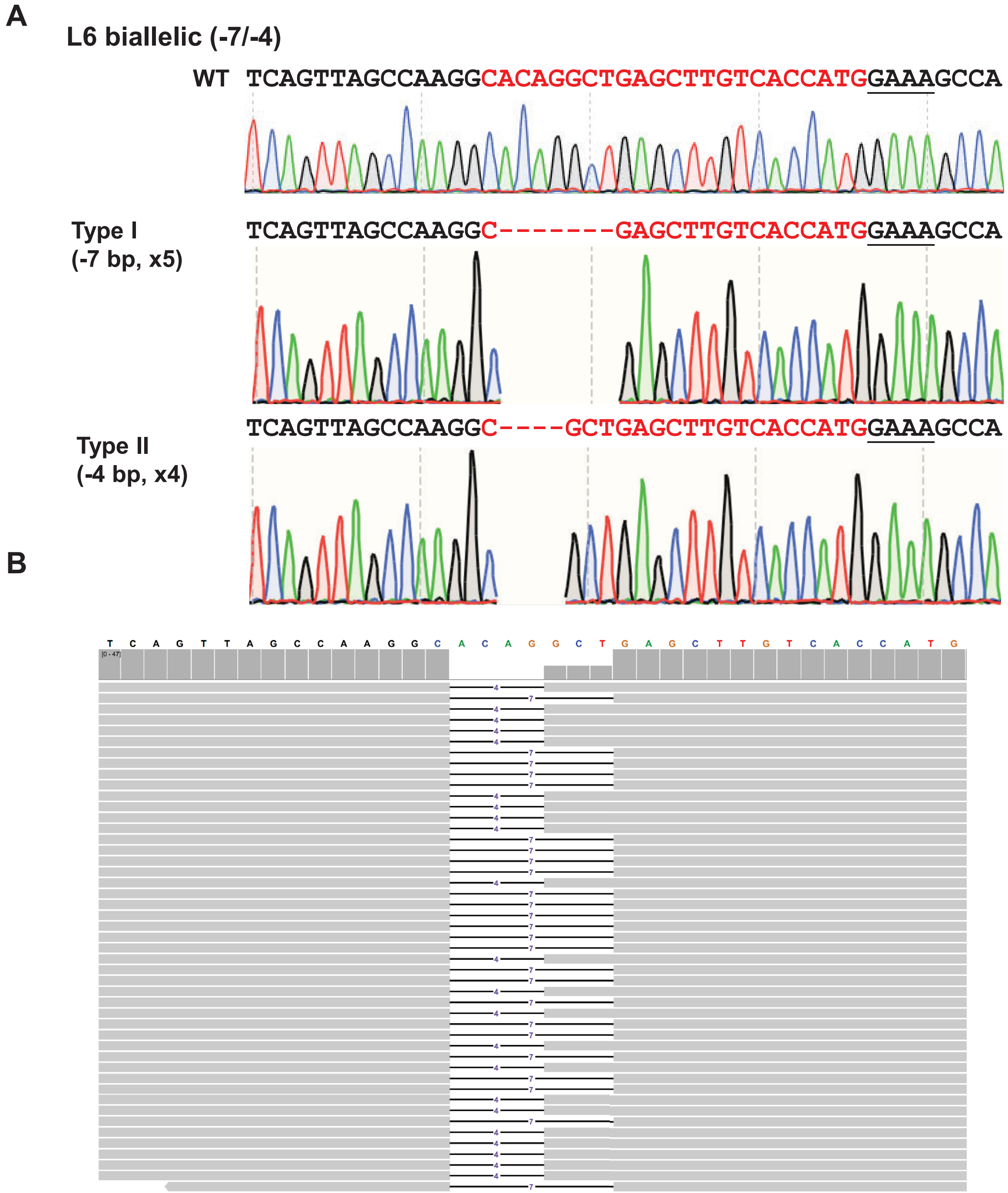

**Extended Data Fig. S8.** Sequencing confirmation of the transgene-free *CsLOB1*-edited *C. sinensis* cv. Hamlin line L6 generated by LbCas12aU/crRNA RNP transformation of embryogenic citrus protoplasts. A. Sequencing confirmation of *CsLOB1* based on PCR amplification and cloning. The representative chromatograms of *CsLOB1* edited *C. sinensis* cv. Hamlin lines. The mutations of both alleles of *CsLOB1* were shown for each line. x indicates number of colonies sequenced. Nucleotide in red indicates crRNA. The underlined GAAA indicates protospacer-adjacent motif (PAM). -: deletion. +: insertion. B. Whole genome sequencing of the edited lines using next generation sequencing. The bases of target site were highlighted by colors other than black. There were two types of deletions of target site, including type I (-7 bp deletion) and Type II (-4 bp deletion), which were shown by the horizontal bar chart. The vertical bar chart showed the sequence depth for each nucleotide.

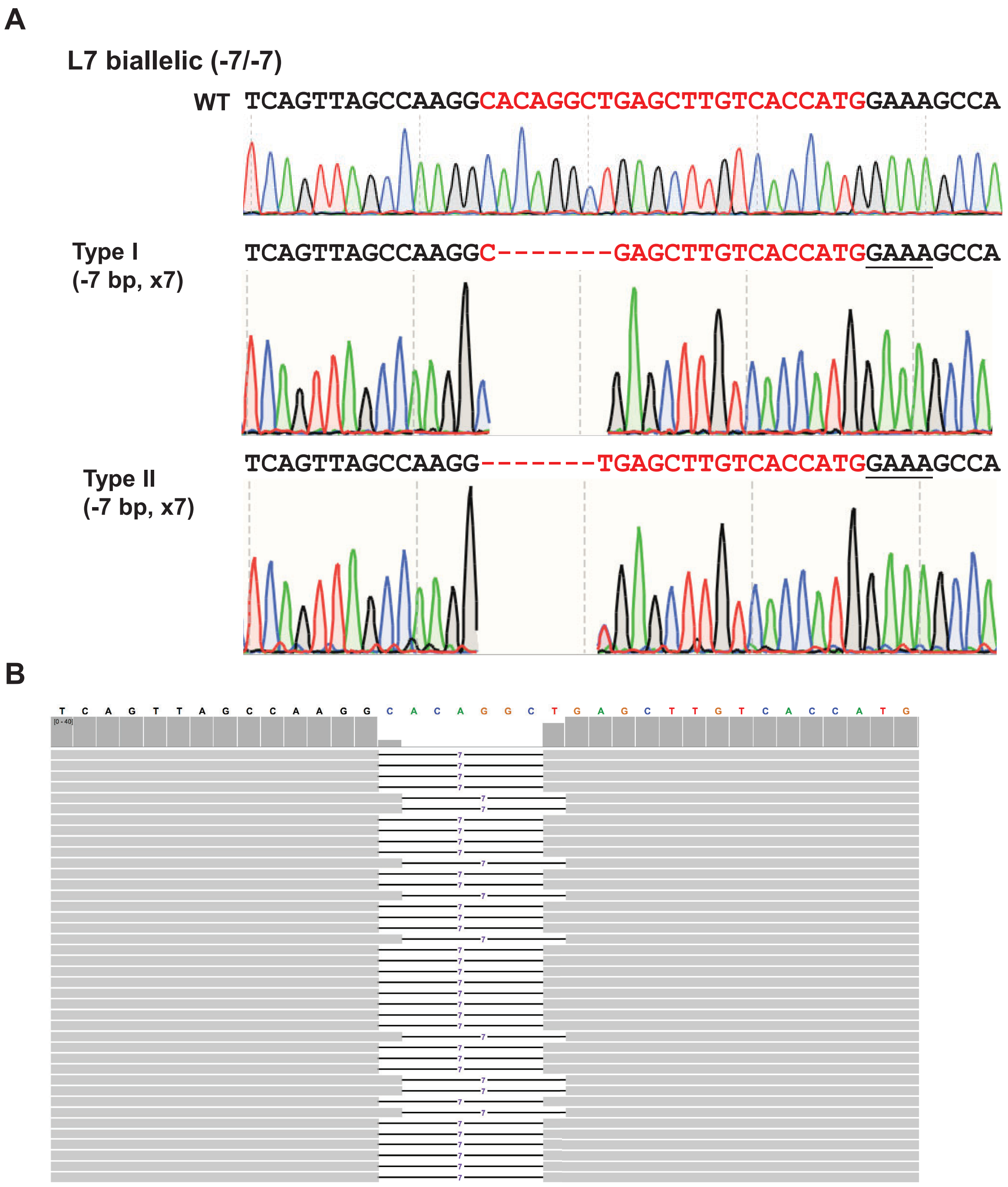

**Extended Data Fig. S9**. Sequencing confirmation of the transgene-free *CsLOB1*-edited *C. sinensis* cv. Hamlin line L7 generated by LbCas12aU/crRNA RNP transformation of embryogenic citrus protoplasts. A. Sequencing confirmation of *CsLOB1* based on PCR amplification and cloning. The representative chromatograms of *CsLOB1* edited *C. sinensis* cv. Hamlin lines. The mutations of both alleles of *CsLOB1* were shown for each line. x indicates number of colonies sequenced. Nucleotide in red indicates crRNA. The underlined GAAA indicates protospacer-adjacent motif (PAM). -: deletion. +: insertion. B. Whole genome sequencing of the edited lines using next generation sequencing. The bases of target site were highlighted by colors other than black. There were two types of 7 bp deletions of target site which were shown by the horizontal bar chart. The vertical bar chart showed the sequence depth for each nucleotide.

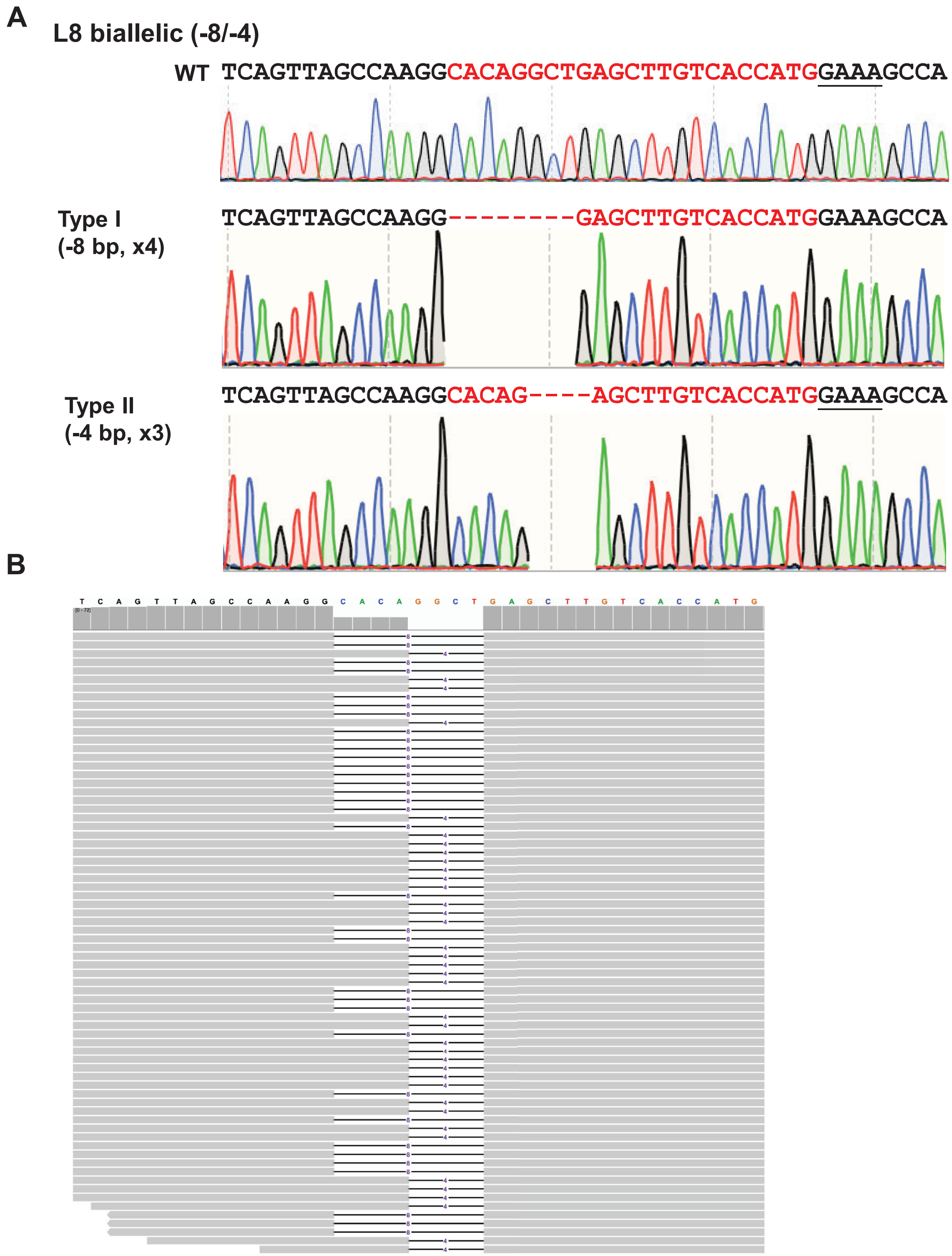

**Extended Data Fig. S10.** Sequencing confirmation of the transgene-free *CsLOB1*-edited *C. sinensis* cv. Hamlin line L8 generated by LbCas12aU/crRNA RNP transformation of embryogenic citrus protoplasts. A. Sequencing confirmation of *CsLOB1* based on PCR amplification and cloning. The representative chromatograms of *CsLOB1* edited *C. sinensis* cv. Hamlin lines. The mutations of both alleles of *CsLOB1* were shown for each line. x indicates number of colonies sequenced. Nucleotide in red indicates crRNA. The underlined GAAA indicates protospacer-adjacent motif (PAM). -: deletion. +: insertion. B. Whole genome sequencing of the edited lines using next generation sequencing. The bases of target site were highlighted by colors other than black. There were two types of deletions of target site, including type I (-8 bp deletion) and Type II (-4 bp deletion), which were shown by the horizontal bar chart. The vertical bar chart showed the sequence depth for each nucleotide.

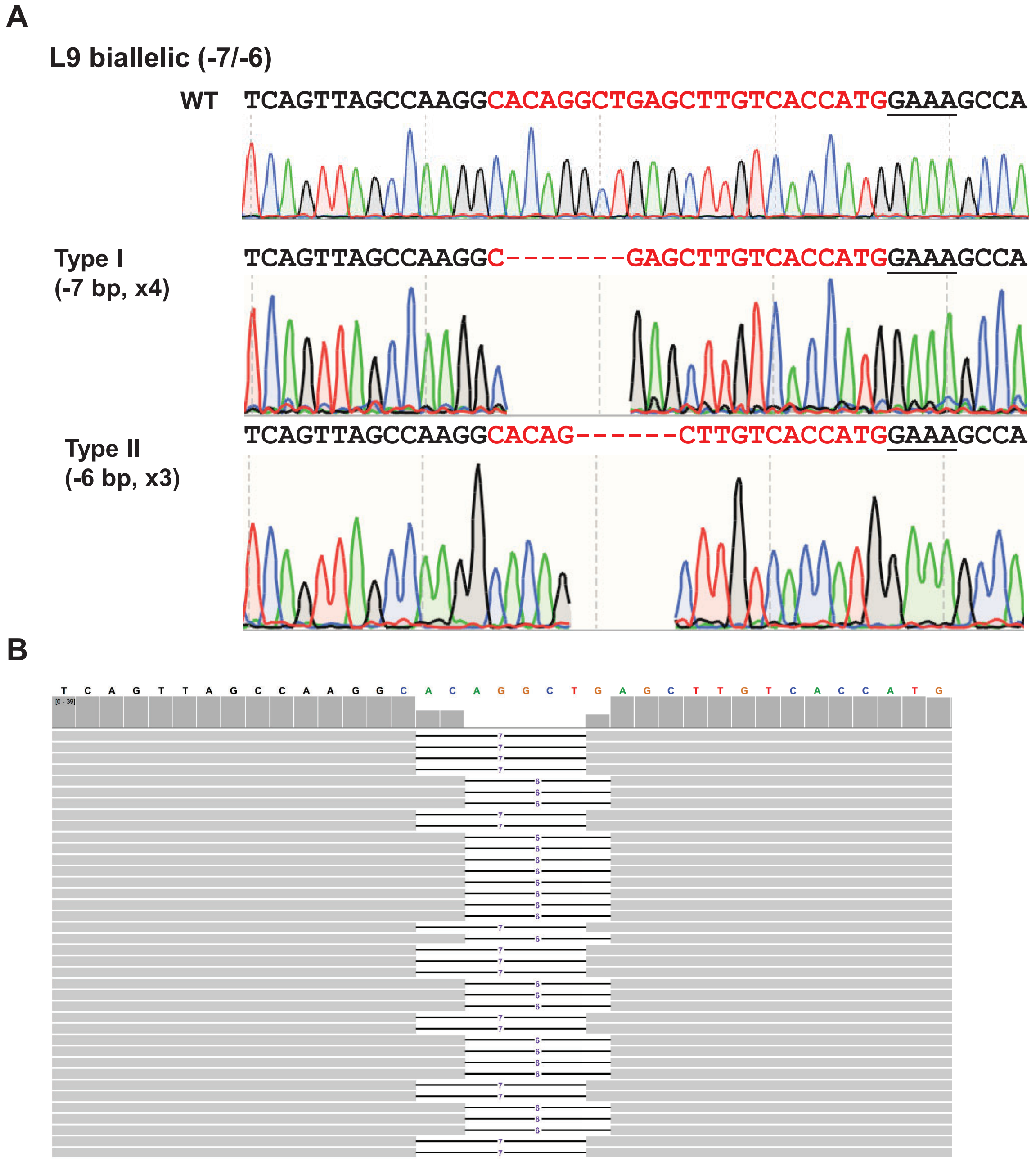

**Extended Data Fig. S11.** Sequencing confirmation of the transgene-free *CsLOB1*-edited *C. sinensis* cv. Hamlin line L9 generated by LbCas12aU/crRNA RNP transformation of embryogenic citrus protoplasts. A. Sequencing confirmation of *CsLOB1* based on PCR amplification and cloning. The representative chromatograms of *CsLOB1* edited *C. sinensis* cv. Hamlin lines. The mutations of both alleles of *CsLOB1* were shown for each line. x indicates number of colonies sequenced. Nucleotide in red indicates crRNA. The underlined GAAA indicates protospacer-adjacent motif (PAM). -: deletion. +: insertion. B. Whole genome sequencing of the edited lines using next generation sequencing. The bases of target site were highlighted by colors other than black. There were two types of deletions of target site, including type I (-7 bp deletion) and Type II (-6 bp deletion), which were shown by the horizontal bar chart. The vertical bar chart showed the sequence depth for each nucleotide.

A

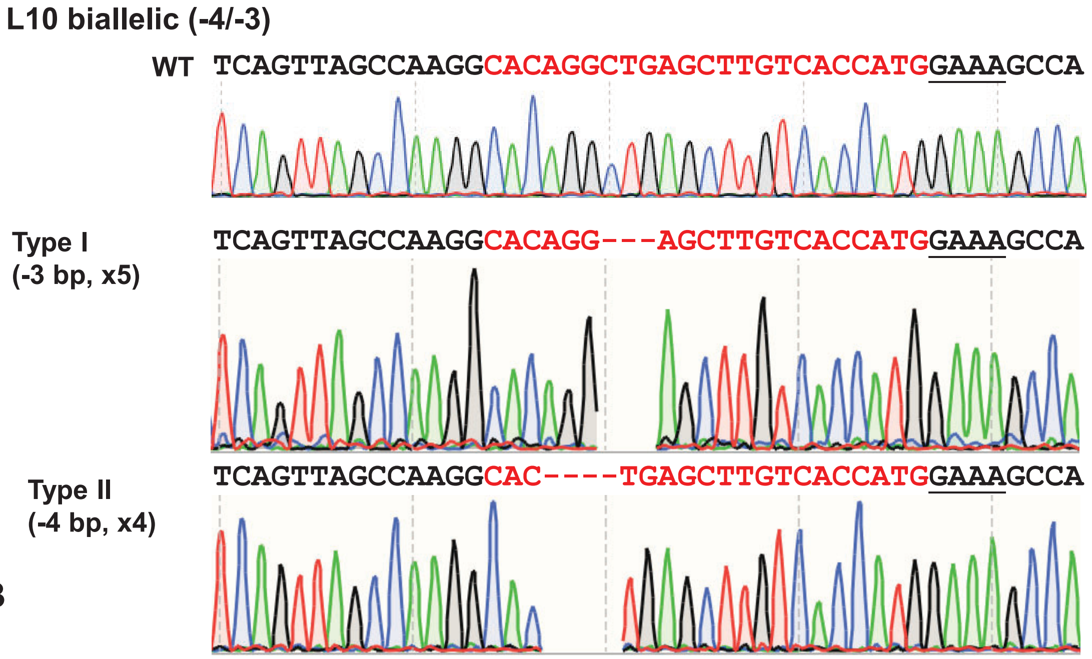

B

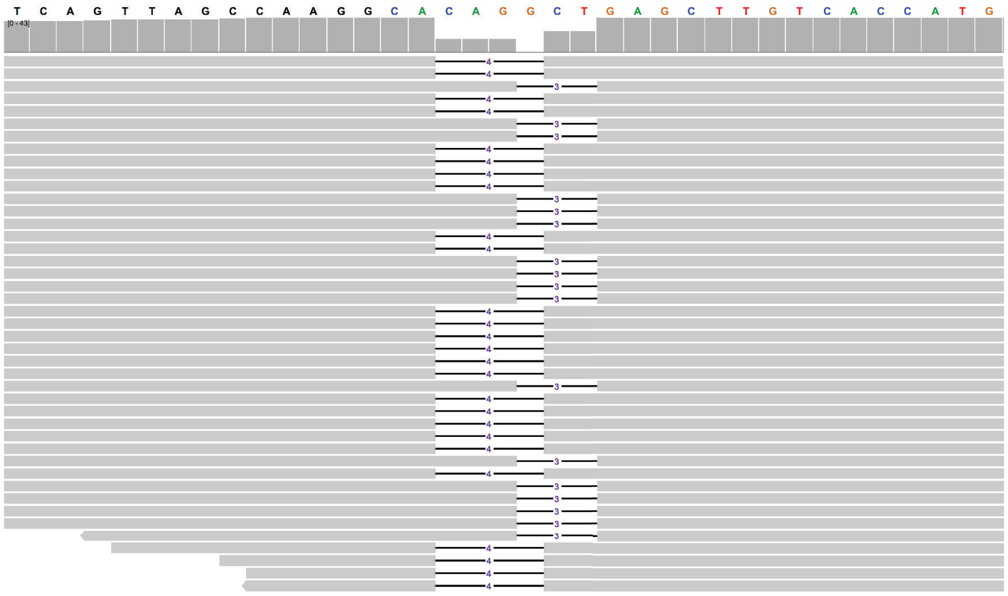

**Extended Data Fig. S12.** Sequencing confirmation of the transgene-free *CsLOB1*-edited *C. sinensis* cv. Hamlin line L10 generated by LbCas12aU/crRNA RNP transformation of embryogenic citrus protoplasts. A. Sequencing confirmation of *CsLOB1* based on PCR amplification and cloning. The representative chromatograms of *CsLOB1* edited *C. sinensis* cv. Hamlin lines. The mutations of both alleles of *CsLOB1* were shown for each line. x indicates number of colonies sequenced. Nucleotide in red indicates crRNA. The underlined GAAA indicates protospacer-adjacent motif (PAM). -: deletion. +: insertion. B. Whole genome sequencing of the edited lines using next generation sequencing. The bases of target site were highlighted by colors other than black. There were two types of deletions of target site, including type I (-3 bp deletion) and Type II (-4 bp deletion), which were shown by the horizontal bar chart. The vertical bar chart showed the sequence depth for each nucleotide.

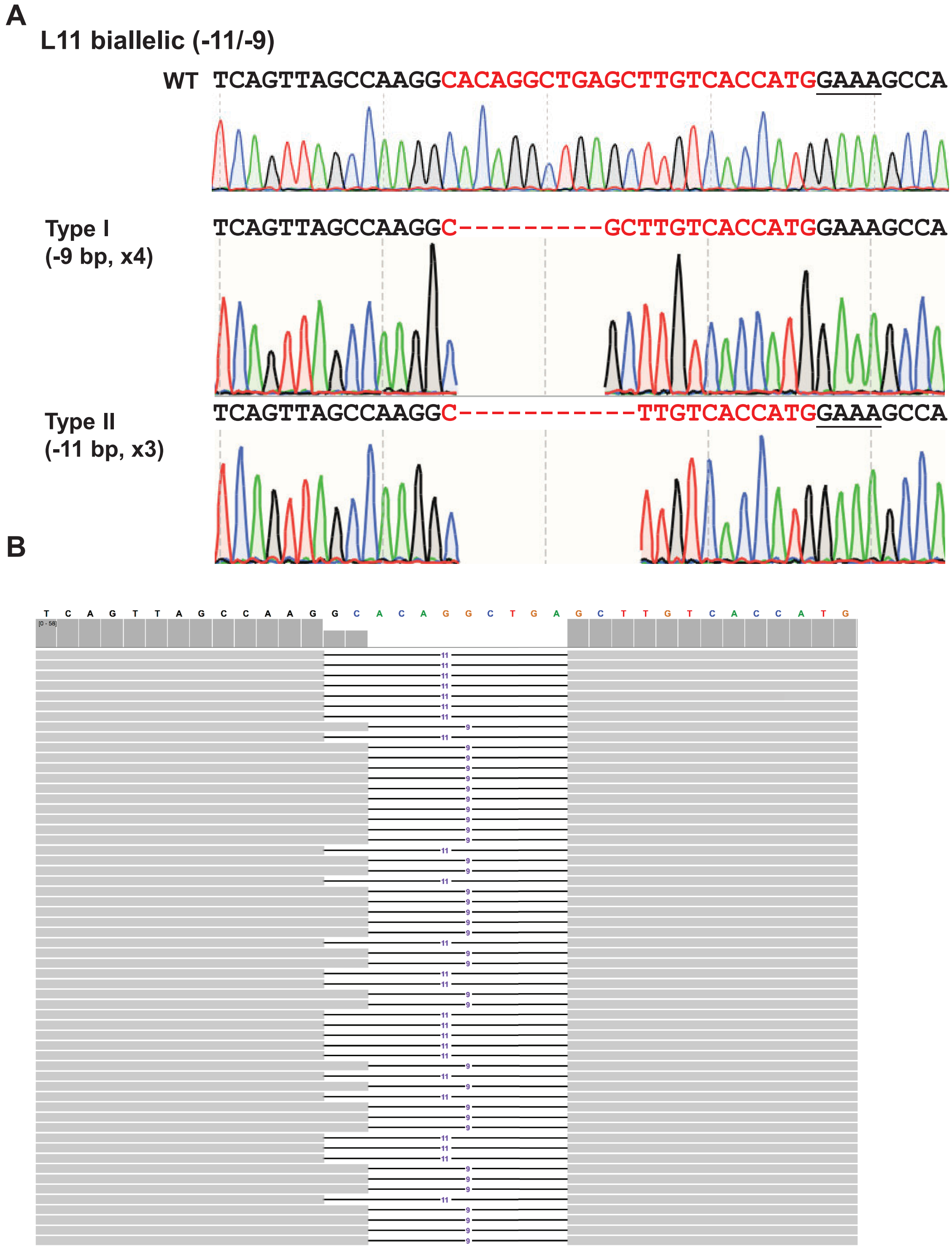

**Extended Data Fig. S13.** Sequencing confirmation of the transgene-free *CsLOB1*-edited *C. sinensis* cv. Hamlin line L11 generated by LbCas12aU/crRNA RNP transformation of embryogenic citrus protoplasts. A. Sequencing confirmation of *CsLOB1* based on PCR amplification and cloning. The representative chromatograms of *CsLOB1* edited *C. sinensis* cv. Hamlin lines. The mutations of both alleles of *CsLOB1* were shown for each line. x indicates number of colonies sequenced. Nucleotide in red indicates crRNA. The underlined GAAA indicates protospacer-adjacent motif (PAM). -: deletion. +: insertion. B. Whole genome sequencing of the edited lines using next generation sequencing. The bases of target site were highlighted by colors other than black. There were two types of deletions of target site, including type I (-9 bp deletion) and Type II (-11 bp deletion), which were shown by the horizontal bar chart. The vertical bar chart showed the sequence depth for each nucleotide.

**A**

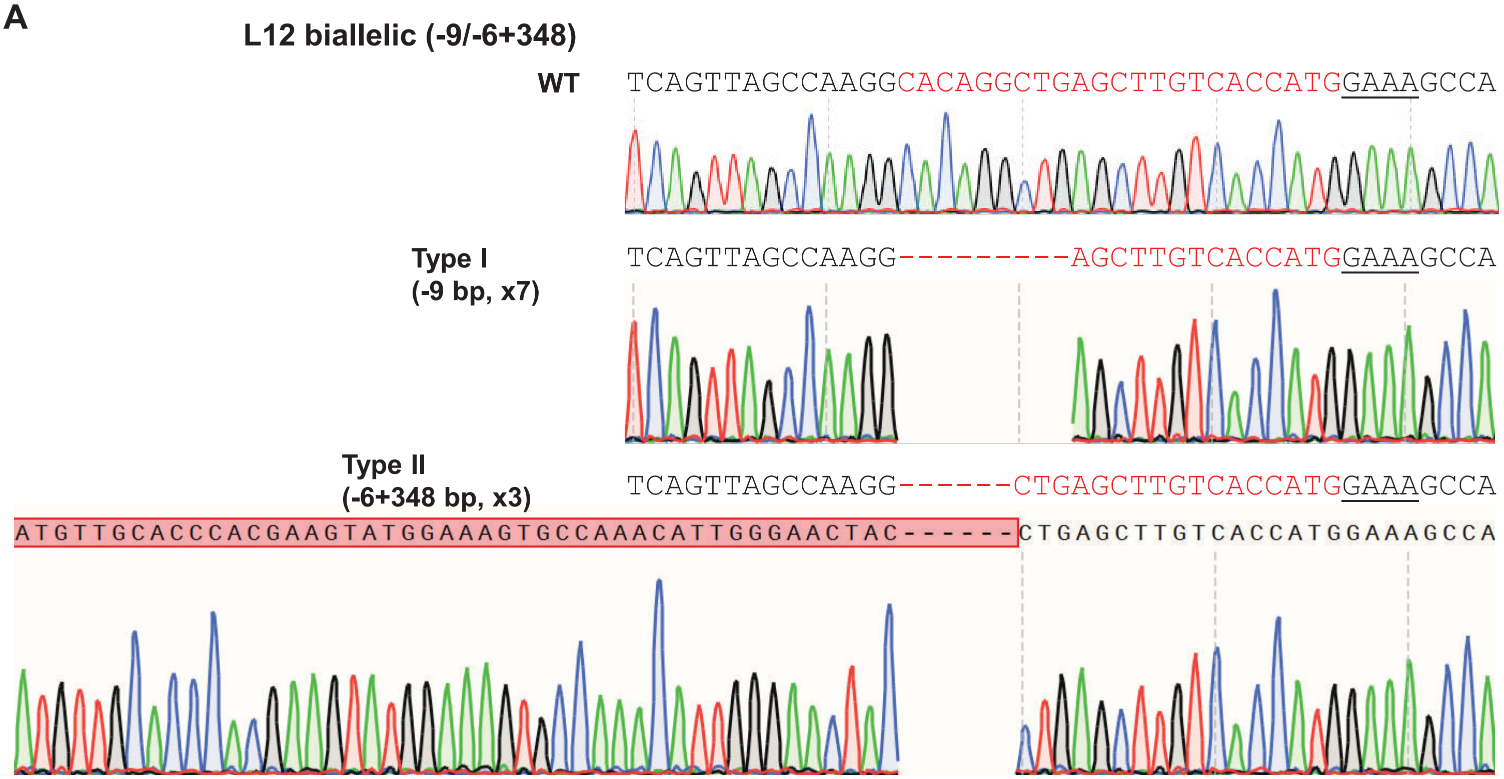

**B**

CCTTGAAAAATTCATATTAACGTTATCAATGATTTTTTTTTAATAGTTTTACCACTTATTTTTTTATAACACCTTGGTAATTTTGACATTAGGTAGCAATATAATACGA
TAAAATTCACCTCCATGTAATTTGAAGTTCTTTTCAATAATTTTTTTGACAAATTTTATAGAAGAATTTAACCTTTTTTTTTTTGGTTCAAACGAAGAAATGTTTCCGT
CATTCAATTAAAATTAATGACATCATCTAGTGGCTCGGTGACATACGCTTTAGATACAATTGTCATTCTTGCCTTTTCCTTTCTCTATATAAACCCCTTTTGCCTTG
AACTTTGTTTCAACTAAAGCAGCTCCTCCTCATCCCTTACTGTCTTTGCTTTCTCACTAACTACTACAACCCAACAGTTTTCTTCTCTCAAAAATGGAATGCAAACA
CAAAATTAATGTAGCAATCCCAATCACTAATATGAAGAACACTCAATTCTCATCTCCATCTACTTTCTCTACTTCTCCTCCTTCTCAATCTTCTCCACGCTTCCCTT
CTCCTAATCATCAACAATTGTCTTCTCCAGAATCTTCTCCAAGCTTTAAAGCTTCTCCTTCACAATCCTCTCCAAATCTTGCAGCTCCCCTCTCTCCGCCGCCTAT
AGTTCTTAGCCCTTGTGCTGCTTGCAAAATCCTCCGCCGCAGATGCGTCGAGAAATGTGTTTTAGCTCCATATTTTCCACCAACCGAACCATACAAGTTCACCAT
TGCTCATAGAGTCTTCGGTGCTAGCAATATCATCAAGTTCTTGCAGGTATGCACTTCTTTTGTATGTGATAAATTCAAACTAATTAAATGTCCAACCATTTTTTTTC
TAATTGGGAGAAAAAAAAAACTTGTTAATTGTTTTATTTTCATCAATTAGTTGTGTGATTAGACTTTGGAGTGGTTGATTGTTCCACTCTTTTTTGGAAACTTACGG
ACTTCTCTAATCAAAAGAAAAGAGAGTGTGACATTTCAACTGATTTCATGCAACTTTAATGTTTGTTTTCAATTCACTTTCTTTAATTTAAATAGAGTAGATTATTCA
AATGGATCCTTTTTATTTCTCCTTATAAAGGAATCATCATCACTCTTCCTTGACGGGGCACTGGCGCAGGGTCAATCTGCCTCGTATACAAGTTTTACAAAGCTT
AAATTTTGCCCCTTTCTAGAAAATATTGTGAATTTTACAAAGTCAAACAATGAGTATGAACTTAGCACATGGGTCCACATTAGAAACCAATTCCATGCCCCAGCAG
CTTAATATCATAAAGTTAGCACATGGCCCGTTAAGACTTAAGCTTTGACAGAATCCAGCGCTCAGAGAACGGTTTAAAATGTTTCTCATTTAGACTAGACTTGCA
AATTAAAATTAAAATATTTCAGTCATTTTATTTTATTAAAATGTAAGGGAGGGTAAATCCCAAATTAATTTTTCGCCTCCCATCTTCTCAAGATTCAAATTCTGGTG
GTCAGACAATAAAATTGTTTGAAAACGATTTGAAGCAGCCTTGGGGCTGAAAATTTCAGTCATCCATATTTGCACACAAACAATGTCTCATGCCATTAAAAATTTT
CCAGGAACTGCCAGAATCTCAACGAGCAGATGCAGTGAGCAGCATGGTCTATGAAGCAAGTGCCAGAATCCGGGATCCTGTTTACGGCTGCGCCGGGGCTAT
TTGCCATCTCCAGAAACAAGTCAGTGAGCTTCAGGCTCAGTTAGCCAAGG**AAATTCTGGGGTGGTCATTATAGCGGTTCCTTCTGATCCTAGAAAACGTCCGAA**
**AAAACCTGCTACGGAACTACCTAGCAGGGGCAAAAATACGATAAGTAGATACATAATTTCGAGTGTGATCAGACAACCAAAAATCAGACAATGAGAGAGCGG**
**CTGAGTGATAGATTGATCGACGTTCGGAGAACGCTCGACCGAAGGAATTTTGCACAAGGTGAATTGTAAGCCCCAACGACTTCGGAACGAGCATTTTCGGGG**
**ACTAGCCCGCTTAGGTTGTAACTTGATCATGAGCGAATTTGTTGATGTTGCACCCACGAAGTATGGAAAGTGCCAAACATTGGGAACTACCTGAGCTTGTCAC**
**CATG**GAAAGCCAGCAACGCAATTTAATAACTCTAATTTGCATGGAAATGGCACAATCTCAAGAACAAGTCTTGCAGCAGCAGCAGCAGCAGCAGCAACAGTTCA
TGGATACTAGCTGTTTTTTGGATGACAATGGTATTGGATCAGCTTGGGAGCCTCTGTGGACATGATCAAGAGAAATTAAAGCAAGATTGTTGAAATTTTAACCTT
TTAAGAGATTATTTACATAAAGCTAAACATACTTAATTATAAAAGTTTCTGATCAATAATTAAAGTTATTTGCTGCGCGGTAGATGGGAGTGATTATTATGTGCTTT
AATTTTCATTAGTCTTGTTGACAAAAAGGAATCTTTGAACCATCTGGAGAAGTCCTTTGTTAACGGTTCGAGATTAATTATTAGTTTATCTTTATTTACATTTAGTG
AAATTTTGTTTTTAACTAATTTTATAGACATAAATAACCCAACCAAGATGGGAATTCAGTGC

Legend: Exon **crRNA** **PAM site** insertion sequence

**Extended Data Fig. S14.** Sequencing confirmation of the transgene-free *CsLOB1*-edited *C. sinensis* cv. Hamlin line L12 generated by LbCas12aU/crRNA RNP transformation of embryogenic citrus protoplasts. A. Sequencing confirmation of *CsLOB1* based on PCR amplification and cloning. The representative chromatograms of *CsLOB1* edited *C. sinensis* cv. Hamlin lines. The mutations of both alleles of *CsLOB1* were shown for each line. x indicates number of colonies sequenced. Nucleotide in red indicates crRNA. The underlined GAAA indicates protospacer-adjacent motif (PAM). -: deletion. +: insertion. For L12 genotype, one allele contained both 6 bp deletion and 348 bp insertion of *C. sinensis* mitochondrial sequence. Only part of the insertion was shown. B. One allele of *CsLOB1* sequence in L12 showing insertion sequence and 6 bp deletion (CACAGG).

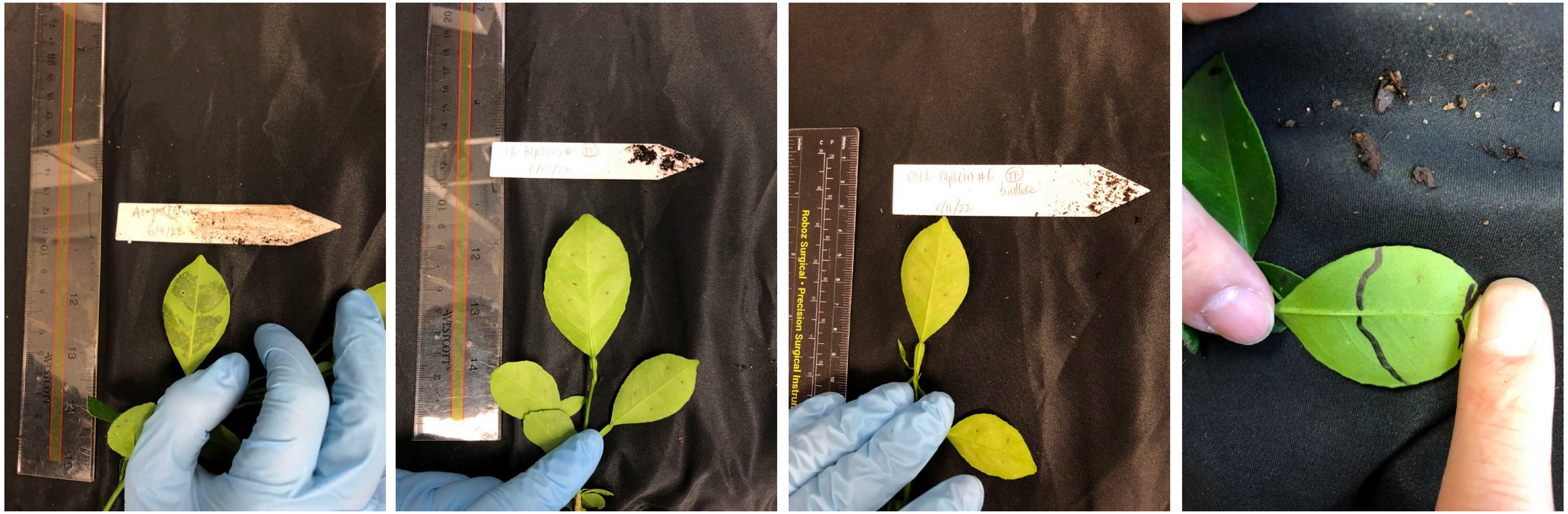

**WT + Xcc** | ***cslob1* + Xcc** | ***cslob1* + Xcc** | ***cslob1* + Xcc**

**Extended Data Fig. S15.** Canker symptoms on wild type *C. sinensis* cv. Hamlin and *cslob1* mutants. Fully expanded citrus leaves were inoculated with Xcc at $10^7$ CFU/ml using needleless syringes. The picture was taken at 9 days after inoculation.

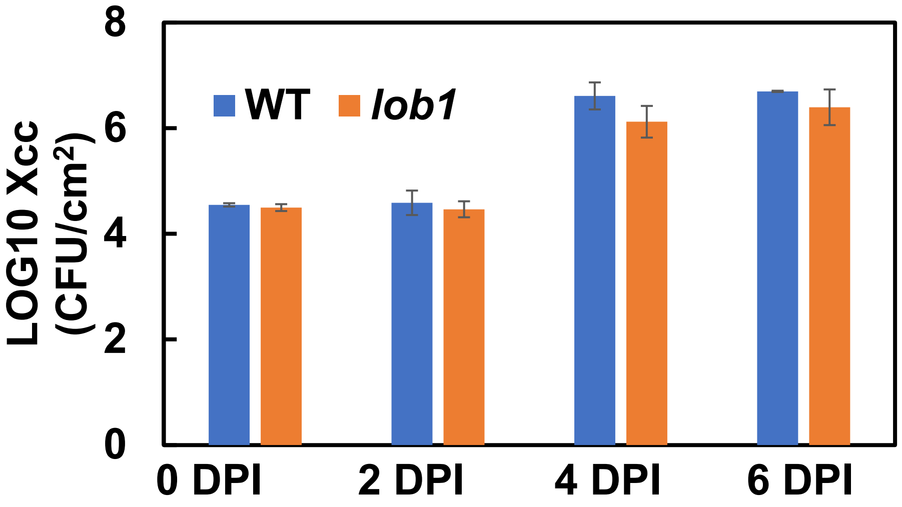

**Extended Data Fig. S16.** *Xanthomonas citri* subsp. citri growth in wild type and transgene-free *cslob1* mutant of *Citrus sinensis* cv. Hamlin. Fully expanded citrus leaves were inoculated with Xcc at $10^7$ CFU/ml using needleless syringes. Three biological replicates were used. DPI: Days post inoculation. 0 DPI indicates right after inoculation.
